# Identification of clinical carbapenemases, 478 novel β-lactamases and putative seven new bacterial phyla from wastewater contaminated high arctic fjord sediments

**DOI:** 10.1101/2024.11.29.625992

**Authors:** Manish P. Victor, Lise Øvreås, Nachiket P. Marathe

## Abstract

Arctic microbiota is enigmatic and highly underexplored. With the aim of understanding the resistome and microbiota of high-Arctic fjord sediments and the effect of wastewater discharge on sediment microbiota, we analysed sediments using metagenomics. We show presence of 888 clinically relevant antibiotic resistance genes including genes coding resistance against last-resort antibiotics such as carbapenems, colistin, vancomycin, linezolid and tigecycline in sediment microbiota. Using computational models, 478 novel β-lactamases belonging to 217 novel β-lactamases families were revealed in sediment microbiota from Adventfjorden in Svalbard. We identified hosts for 69 novel gene families and showed that these genes are widespread in the Arctic environment. Further, in 644 metagenome-assembled genomes (MAGs) from sediment metagenomes >97% belong to novel taxa, representing seven putative novel phyla. These MAGs encoded important functions like nutrient cycling and methane metabolism etc. Our study demonstrated mixing of human associated bacteria and Arctic sediment microbiota. It provides the first comprehensive dataset of the distribution and diversity of novel microbes and β-lactamases in the high Arctic sediments.

## Introduction

The Polar Regions are geographically distant ecotypes on Earth with extreme environmental conditions like low temperature, low nutrient availability and highly variable light cycles. Microbes are one of the major contributors to total ecosystem biomass, biodiversity and biogeochemical nutrient cycling. Although climate change and human impact has severe effects on the Arctic, little is known about its impacts on Arctic microbial communities (Cavicchioli et al. 2019, Verde et al. 2016). Arctic sediments still represent huge diversity of novel and hitherto uncharacterized microbes. Understanding the diversity and role of microbes in the Arctic is thus of utmost importance for designing mitigation strategies for preservation of the polar environments(Kleinteich et al. 2017, Edwards et al. 2020). The Advent fjord in Svalbard is exposed to human activities like wastewater discharge and increasing tourism. The aim of this study was to understand the impact of anthropogenic activities with respect to microbiota and antimicrobial resistance (AMR) by characterizing microbiota, genomes of novel bacteria and resistome of the Arctic sediment microbiota. Further we aimed at characterizing novel β-lactamases present in the microbiota as well as identifying their potential hosts.

## Results and Discussions

We collected and sequenced sediments from nine different sites in Adventfjorden in duplicates generating a total of ∼450 GB of sequences (SRA accession: SRR28980414-31). The sediment microbiota was dominated by *Psuedomonadota* (45-55% of the total 16S rDNA reads), followed by *Bacteroidota* (10-21%). Around 9 % of the 16S rDNA reads could not be assigned to any phylum and around 8% could not be classified beyond phylum level, thus representing a huge diversity of novel bacteria present in the Arctic sediments (Fig.1a). We did observe variation in microbiota between the sampling sites suggesting micro niches present in high Arctic Fjord sediments. With the aim of characterizing these novel bacteria, we were able to assemble 644 mid-quality metagenome assembled genomes (MAGs). The taxonomy of these 644 MAGs was consistent with the 16S rDNA gene-based microbiota analysis, with *Psuedomonadota* (50%) representing most of the genomes followed by *Bacteroidota* (17%). Five of the 644 genomes belonged to Archaea with one genome representing a novel phylum (Supplementary table 1). Only 23 MAGs could be classified to either genus level (n = 16) or species level (n = 7), thus demonstrating the high abundance of novel taxa in the Arctic sediments (Fig.1b). Around 14% (n = 89) of MAGs were classified into seven putative novel phyla. (Figure 1b, Supplementary fig.1). The metabolic functions deciphered from these genomes show that they perform active roles in nutrient cycling and methane metabolism in the Arctic sediments and thus represent important members of the Arctic microbiota (Supplementary table 2).

**Figure 1.**
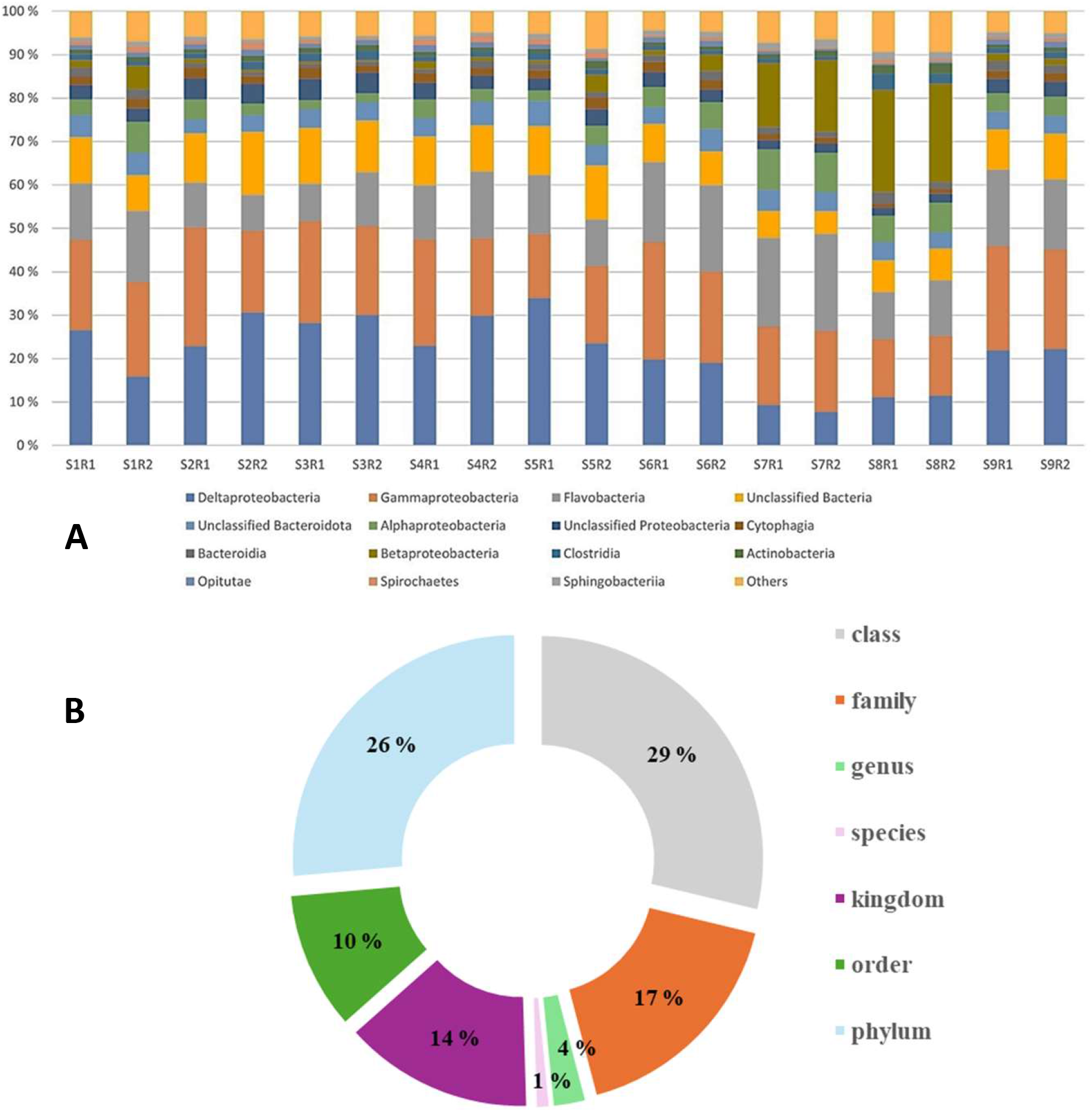
a) Relative abundance of different phyla in the sediment samples. Legend: S-sampling sites and R-replicates eg. S1R1-site 1 replicate 1. b) Taxonomic classification of 639 MAGs (excluding Archea) at the lowest taxonomic unit possible.

To understand the anthropogenic impact on the Arctic microbiota we analysed the abundance of known antibiotic resistance genes (ARGs) (Marathe et al. 2019). We detected a total of 888 different ARGs in these samples encompassing clinically important genes conferring resistance against last resort antibiotics, including carbapenemases NDM, VIM, IMP, KPC and OXA-181, mobile colistin resistance genes *mcr1* and *mcr2*, tigecycline resistance gene *tet*(X) as well as linezolid resistance genes *poxtA, cfr and optrA* (Supplementary table 3). Suggesting the presence of human associated bacteria, most likely pathogens, in the sediment microbiota owing to continuous wastewater discharge into the fjord (Marathe et al. 2019, Zhang et al. 2021). Most of the clinically relevant ARGs have originated in the environmental bacteria, it serves as a source for acquiring new ARGs for pathogens (Forsberg et al. 2014). The presence of human associated bacteria in the Arctic coupled with climate change, make the genetic exchange between Arctic microbiota and human pathogens a likely scenario (Edwards et al. 2020) . With high abundance of novel microbes present in the Arctic sediments, we investigated the novel β-lactamases in the sediment microbiota using computational models (Berglund et al. 2019). We discovered 478 unique novel β-lactamases belonging to class A (n = 52), subclass B1/B2 (n = 79), subclass B3 (n = 115) and class D (n = 232), respectively (supplementary table 4, supplementary figure 2). Using gene sequences from β-lactamase database (BLDB) and previously published novel β-lactamases we classified the genes into gene families with cutoff of amino acid identity of 70% (Berglund et al. 2019). We report a total of 217 novel β-lactamase families with 27, 49, 45 and 96 novel families belonging to class A, subclass B1/2, subclass B3 and class D respectively (supplementary table 4, Supplementary figure 3a-d). Our study thus almost doubles the number of β-lactamase families (BLFs) present in BLDB. Class B (subclass B1, B2 and B3) are clinically very important and represent carbapenem-hydrolyzing metallo-β-lactamases (MBLs) that utilize zinc in the active site (Boyd et al. 2020). The presence of 94 novel families of MBLs in Arctic microbiota thus suggests the potential of Arctic microbiota as a source of clinically important ARGs for pathogens.

**Figure 2.**
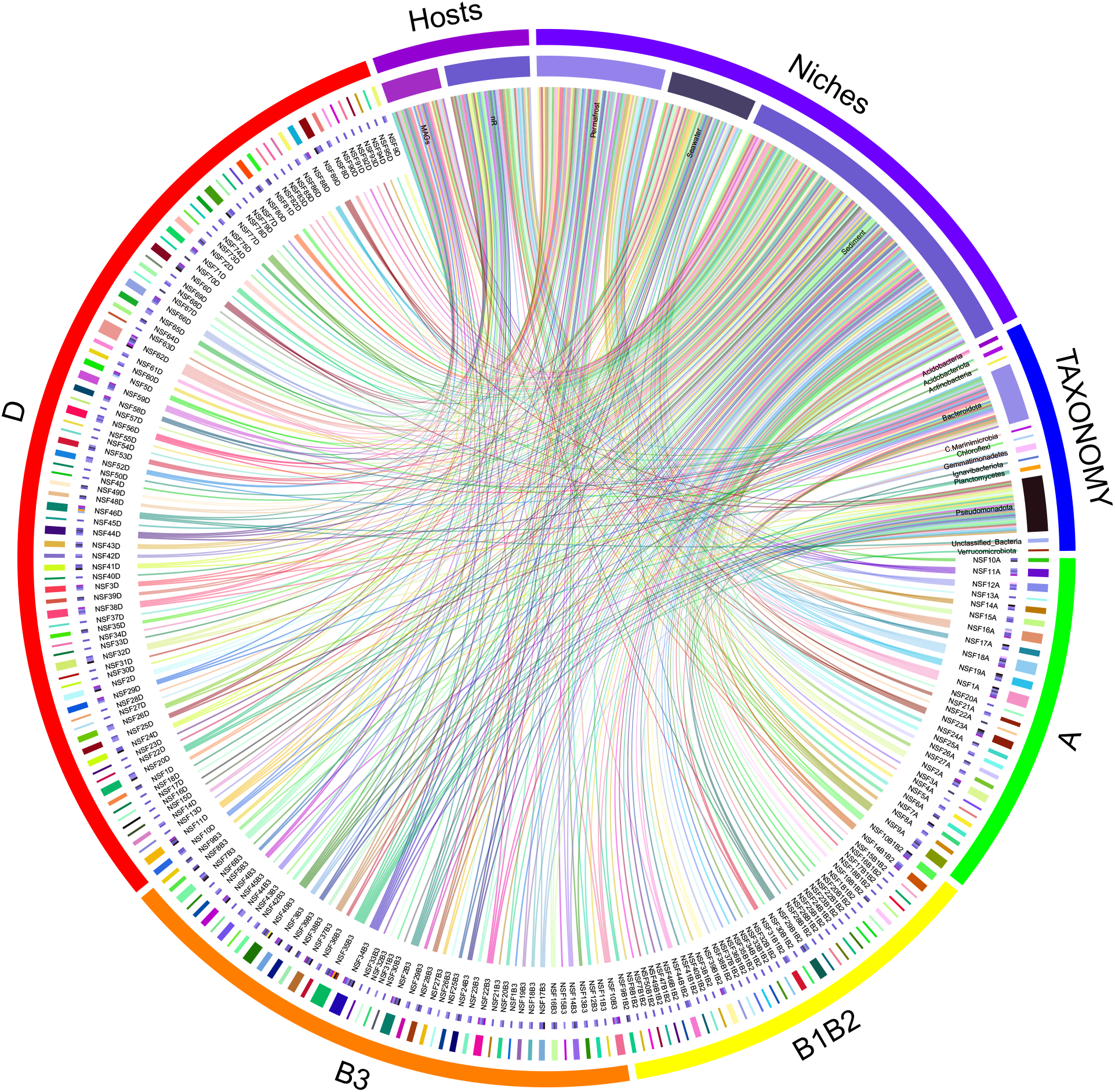
Hosts and niches of novel β-lactamase families detected in our study. NSF denotes novel β-lactamase families. The Circos representation of the presence of β-lactamases across niches (sediment, seawater, permafrost samples) as well as taxonomy of the host species and the source (nR database or MAGs). Links are drawn when a certain gene is found in a certain sample and/or in certain host species representing either MAGs or nR database. Different classes of β-lactamases are represented as A, B1B2, B3, D.

Consistent with differences in microbial community compositions we detected differences in the number of novel ARGs assembled from 9 sites in the Advent fjord. Although, most of the gene families were detected, no full-length β-lactamase was assembled from sampling site 7. All 217 novel gene families were present with varying abundances in the sediments combined, with some families found to be absent in some of the samples (Supplementary figure4).

To identify the potential hosts for the novel BLFs we performed BLAST analysis using a combination of NCBI nR database and MAGs from our study. Hosts for 58 novel BLFs were identified using nR database with 16 families in class A, 10 in subclass B1/2, 11 in B3 and 21 in class D, while MAGs carried a total of 23 novel families, 12 families overlapped between nR database and MAGs. We have thus successfully identified host species for 69 novel BLFs (Fig. 2, Supplementary table 5a-c). Based on metadata all the host organisms represent common marine bacteria, with majority of novel gene families detected in the phylum *Bacteroidota* (34) followed by *Pseudomonadota* (28). Thus, supporting the notion that less-abundant microbes may play a dominant role in the different ecological functions in a given environment (Jousset et al. 2011).

To understand the presence and abundance of novel BLFs in different niches from Svalbard, we used previously published metagenomes from sediments, water and permafrost samples (Zhang et al. 2022, Xue et al. 2020). A total of 168 gene families were detected in sediments from Kongs fjord, while only 65 from permafrost and 46 from seawater metagenomes respectively (Supplementary figure 5). Our data suggests that many of these novel β-lactamases are prevalent in high Arctic niches, while some may be more niche specific.

## Conclusions

Our study provides the most comprehensive analyses of novel microbes and novel β-lactamases present in the Arctic sediments. It demonstrates the effect of mixing novel indigenous microbes and human associated microbiota in the Arctic sediments, and further demonstrates presence of clinically important ARGs and novel ARGs in the Arctic microbiota. With a warming Arctic and continuing anthropogenic impact, Arctic microbiota may serve as a source for emergence of superbug, thus challenging public health issues in the future.

## Methods

### Sample Collection, sequencing, quality filtering and taxonomy

#### Sample Collection

Sediment samples were collected from nine sites in Adventfjorden in Svalbard in duplicates using a sediment grab and a boat (Supplementary figure2). The samples were collected in sterile 50ml tubes, frozen and transported to the laboratory.

##### 1b. DNA extraction and sequencing

The total DNA was extracted from the sediment samples using DNeasy PowerSoil kit (Qiagen Germany), following the manufacturer’s protocol. The extracted DNA was quantified, using QubitTM 2000 Spectrophotometer (Thermo Scientific, USA). Sequencing libraries were prepared, using Nextera Library Prep Kit (Illumina, USA), with 2×150bp chemistry, at Norwegian Sequencing Centre, Oslo.

#### Quality filtering of the reads

The obtained FASTQ reads were quality trimmed with TrimGalore (Version 0.6.10) which is a wrapper around Cutadapt (Version 2.6) and FastQC (Version 0.12.1). (trim_galore --quality 30 – phred33 --fastqc --trim-n –cores {} –paired {Forward_Sequence} {Reverse_Sequence}). The cut-off quality score was set at 30 under the quality scoring schema ASCII+33 i.e. phred33. Self-detection mode for recognising and removing the adapter sequences were used (https://github.com/FelixKrueger/TrimGalore). Stringency was set to 1(Very high) i.e. a single base-pair overlap from the 3’ end was trimmed and the maximum allowed error was 0.1. The N’s from either side of the reads were also removed. The number of reads obtained in each sample is presented in Table M1.

#### Taxonomic classification based on SSU

Metaxa2(V 2.2.3) (Bengtsson-Palme et al. 2015) pipeline was used for the taxonomic assignment of the trimmed reads using small subunit ribosomal ribonucleic acid. The basis of the Metaxa2 SSU database is the SILVA2 reference release 111 and Mtozoa3 release 10. Metaxa2 -o metaxa_out -1 {Forward Sequence} -2 {Reverse Sequence} -f fastq -g ssu –mode metagenome –usearch v11.0.67_i86linux64 –usearch_bin usearch –cpu {number of cores} –table T –reltax T. In the next step we have used the metaxa taxonomy traversal tool on the output file obtained from the previous step as the input for : metaxa2_ttt -i {out_taxonomy.txt} -o {outfile} for filtering out taxonomy at the different taxa levels .i.e.(Phylum-Species).

### Detection of metal, biocide, anti-microbial resistance resistance genes, mobile genetic elements and virulence factor detection from quality reads

The detection of different genes was performed as described in (Marathe et al. 2017; Berglund et al. 2019) with the following modifications. Antimicrobial resistance genes local database was prepared by combining datasets from NCBI AMRFinderPlus(Feldgarden et al. 2021), CARD(Jia et al. 2017), AG-ANNOT(Gupta et al. 2014). Antibacterial biocide and metal resistance genes were obtained from BacMet(Pal et al. 2014). Sequences for mobile genetic elements10 were obtained from (Pärnänen et al. 2018). Then the quality trimmed reads were analysed for the respective genes using Usearch v11.0.667_i86linux64 (Edgar 2010). usearch -userach_global {forward.fastq} -db {local_database} -evalue 0.00001 -id 0.9 -query_cov 1 -blast6out {out_file} -strand positive -maxaccepts 1-threads{number of threads}.

We used query coverage of 100% and percentage identity of 90% for the trimmed reads with the different datasets. An in-house python script was written to count the hits of trimmed-reads that matched with genes in the different datasets. The reads were normalized to the length of the respective genes and were further normalized with length normalized 16S rRNA gene copies, expressing abundance as gene copies/16S rRNA copy. The data is plotted as a histogram showing the varied abundance of different genes.

### Assembly and characterization of β-lactamases

fARGene (Berglund et al. 2019) (Fragmented Antibiotic Resistance Gene identifier) (Version v.0.1) is a tool which takes fragmented metagenomic sequences as in-file and produces a predicted full length antibiotic resistance genes. The pipeline utilizes hidden Markov model (HMM) based upon different resistance genes. Present study utilizes Class A β-lactamases, Subclass B1 and B2 βlactamases, Subclass B3 β-lactamases, Class C and Class D β-lactamases optimized models. Command line (fargene --infiles {Forward_Sequence} {Reverse_Sequence} --hmm-model {model_name} --score None --meta --output {folder_out} --force --processes {cpu} –rerun -- amino-dir class_a_out/tmpdir/ --orf-finder) has been used.

### Clustering known and new β-lactamase families (BLFs)

In-order to group the assembled β-lactamase and to identify the novel BLFs clustering was carried using USEARCH. Initially for each of the aforementioned classes the known gene sequences were collected from β-Lactamase DataBase (BLDB) (Naas et al. 2017) and (Berglund et al. 2017). Thereafter, under each class of the BL the amino-acid sequences were merged resulting in all the sequences in four fasta files (Classes A, B1/B2, B3, D). In the next step redundant sequences were removed by clustering at 100% identity using usearch -cluster_fast {in_file.fasta} -id 1 -centroids {unique_file.fasta}. In the following step clustering was done at an identity of 70% for the unique files as suggested in (Berglund et al. 2019) for identifying novel β-lactamses families using usearch -cluster_fast {unique_file.fasta} -id 0.7 -clusters cluster_dir/c_. With this step under each class we obtained many cluster fasta files (CFF) named as (c_0,c_1,c_2……) showing the clustering of different sequences in each CFF. The An in-house python script was written to :

(i) extract one sequence from the CFFs which exclusively have β-lactamases obtained from our analysis. These filtered sequences represent novel β-lactamase families and taken as representative of the cluster.
(ii) Wherever the program encounters clusters exclusively with known β-lactamases it extracts one sequence as the representative of the cluster.
(iii) Wherever the program encounters clusters with known and novel β-lactamases it extracts known β-lactamases and taken as representative of the cluster.
(iv) For each cluster the number of ARGs in them have been recorded and represented in the phylogenetic trees with the representative sequences of the cluster.

### Phylogeny of β-lactamse

All of the composite β-lactamases resulted above in each class was aligned with MAFFT (Version v7.505) (Katoh 2002) and a newick tree file was generated with FASTTREE (Version 2.1.11) (Price, Dehal, and Arkin 2010). The phylogenetic trees were reconstructed with treedyn in phylogeny.fr (Dereeper et al. 2008).

### Assembly of MAGs

Assembly was done with SqueezeMeta (V1.6.2)(Tamames and Puente-Sánchez 2019). It is a fully automatic pipeline for metagenomics/meta transcriptomics covering all steps of the analysis. Assembly for the quality trimmed reads fastq files was done using Megahit (Li et al. 2015). Contigs (<1000 bps) were removed and contig statistics were done using prinseq (Schmieder and Edwards 2011). RNAs were predicted using Barrnap (V 0.9)(Wang et al. 2007). 16S rRNA sequences were taxonomically classified using the RDP classifier (Wang and Cole 2024). Binning was done using Metabat2 (V2.12.1) (Kang et al. 2019). Bin statistics were computed using CheckM (Parks et al. 2015). In the end the MAGs obtained were filtered based upon completeness ≥ 50% and contamination ≤ 10% as suggested by (Benoit et al. 2024).

### Pathway prediction for MAGs

The metabolic pathways have been predicted using a python(Version 2.7.15) based pipeline Fun4me (Sharifi and Ye 2017), executed by python fun4me.py -i {in_file.fasta} -o {outfile}.The pipeline utilizes the funtions of : FragGeneScan (V1.31) (Rho, Tang, and Ye 2010), that predicts the genes in genomes in prokaryotes; RAPsearch (V2.24)(Ye, Choi, and Tang 2011) for rapid protein similarity search tool and MinPath (V1.5) (Ye and Doak 2009) i.e. minimal set of pathways, which is based upon parsimony search for biological pathways determination and reconstruction utilizing the protein family predictions. It helps to achieve conservative but trustworthy biological pathways estimations from the query datasets.

The core datasets of the pipeline are EC2GO(The Gene Ontology Consortium 2019) (https://www.ebi.ac.uk/GOA/EC2GO) , enzyme nomenclature database from the Swiss institute of Bioninformatics (Bairoch 2000), gene ontology annotation database(Gene Ontology Consortium 2004), metacyc(Caspi et al. 2020) pathways data and UniRef90 (Suzek et al. 2015) datasets. The tool generates an HTML file with hyperlinks to MetaCyc and BRENDA (Schomburg et al. 2017) for further understanding the pathways and the enzymes involved in the respective pathways.

### Core-Genome analysis of the MAGs

The genomic relatedness of the obtained MAGs was done using the core genes extracted by the UBCG pipeline(V2) (Na et al. 2018) using default parameters and a maximum-likelihood tree was constructed. First the core genes were extracted for all the 644 MAGs using java -jar UBCG.jar extract -bcg_dir core_genome_dir_name -i input_scaffold.fasta -label output_name then all the obtained core genes were aligned with java -jar UBCG.jar align -bcg_dir {core_genome_dir_name} -prefix {out_name}. Finally, a newick file is generated with the name out_name.UBCG_gsi(92).codon.50.label.nwk which is used to reconstruct the phylogenetic tree as done above.

### Taxonomic classification of unclassified MAGs

Eighty-nine MAGs belonging to unidentified bacteria were recognized. For identifying the most appropriate taxon of these bacteria we again used the core-genome analysis. Initially, the entire NCBI’s dataset for bacteria was downloaded and afterwards an in-house python script was written for extracting the representative bacteria of each validly published genus from the dataset. The scaffold for the respective filtered bacteria was downloaded using NCBI’s esearch -db assembly - query {scaffold_id} | elink -target nuccore | efetch -format fasta > {scaffold_name}. UBCG was used on a combined datasets of MAGs and scaffolds. Finally, a phylogenetic tree was reconstructedexhibiting the clustering of the 89 MAGs with different genomes representing 3540 bacterial genera (Supplementary Method Table 1).

### Identification of putative hosts for the novel β-lactamases families

For identification of novel β-lactamase families (NSF) with known ARGs we have used nR database (27 Nov 2022) and DIAMOND (Buchfink, Reuter, and Drost 2021).The sequences of BLFs were tested against nR database. In the first step diamond blastp --threads {} --db nr.dmnd -- out {out_file}_max1 --header simple --evalue 0.00001 --max-target-seqs 1 --query all_svalbard_fargene.fasta --query-cover 95 --outfmt 6 qseqid qlen sallseqid slen qstart sstart qend send qseq full_qseq full_sseq evalue bitscore length pident stitle salltitles qstrand was used for obtaining the results. Next, we filtered for the unique results with percentage identity *** >=70% under each matching NSF. For each of the matched data the metadata in the NCBI was checked for the habitats from which the bacteria were isolated. Similarly, NSF sequences were checked against MAGs obtained in our study to identify putative hosts for the NSFs.

### Distribution of NSF in different niches and statistical analysis

For understanding the distribution of NSF in different niches we used previously published available sequence data from Svalbard for permafrost (Xue et al. 2020), seawater (T. Zhang et al. 2022a) and Sediment (T. Zhang et al. 2022b) using parameters described in section 2. A heatmap describing the abundance of NSF in different niches was prepared. Finally, a circos plot was made for describing 217 NSFs under all the β-lactamase classes of A, B1B2, B3 and D with different datasets of nR, MAGs, Sediments, Seawater, Permafrost and the various taxons of bacteria.

## Acknowledgements

We acknowledge Institute of Marine research (project number 15930) and the Research council of Norway for funding (Res-Marine project grant number 315266) for funding this work. We thank Nadja Junghardt at IMR and University of Bergen for her help with sample collection.

## Data availability

The short reads are deposited at NCBI SRA under accession numbers SRR28980414-31. The amino acid sequences for assembled beta lactamases are presented in supplementary table 4.

## Declaration

The authors declare no conflict of interest exists.

## Authors contribution

Nachiket Marathe designed the study and acquired the funding. Lise Øvreås and Manish Victor collected samples and carried out the experiment. Nachiket Marathe and Manish Victor analyzed the data. Manish Victor, Nachiket Marathe and Lise Øvreås prepared the draft and revised it. All authors have read the final manuscript and approved it for publication.

## Supplementary materials

**Supplementary table 1**: List of MAGs assembled in our study and their taxonomy.

**Supplementary table 2**: Metabolic pathways detected in 644 MAGs. The results have hyperlink to the metacyc pathways for the pathways and BRENDA for enzymes used in the pathways.

**Supplementary table 3**: Abundance of ARGs, BRGs, metal resistance genes, MGEs and virulence factors detected in different samples.

**Supplementary table 4**: List and amino acid sequences of β-lactamases assembled form the sediment samples.

**Supplementary table 5**: a) List of putative hosts for novel β-lactamase families detected using nR database. b) List of MAGs carrying novel β-lactamase families. c) List of novel β-lactamase families detected for which hosts were detected in nR database and MAGs.

**Supplementary figure 1**: Core-genome phylogenetic analysis for the representative species of validly published genera and 89 MAGs from our study that could not be assigned to any phylum.

**Supplementary figure 2**: Sampling sites co-ordinates and the number of β-lactamases detected in each site.

**Supplementary figure 3**: a) Phylogenetic tree of class A β-lactamases. b) Phylogenetic tree of subclass B1B2 β-lactamases. c) Phylogenetic tree of subclass B3 β-lactamases. d) Phylogenetic tree of class D β-lactamases.

**Supplementary figure 4**: Abundances of 217 novel β-lactamase families across sampling sites in our study.

**Supplementary figure 5**: Presence and abundance of novel β-lactamase families in sediment, permafrost and seawater samples from Svalbard.

