## Supplementary 2 for "Identification of clinical carbapenemases, 478 novel β-lactamases and putative seven new bacterial phyla from wastewater contaminated high arctic fjord sediments"

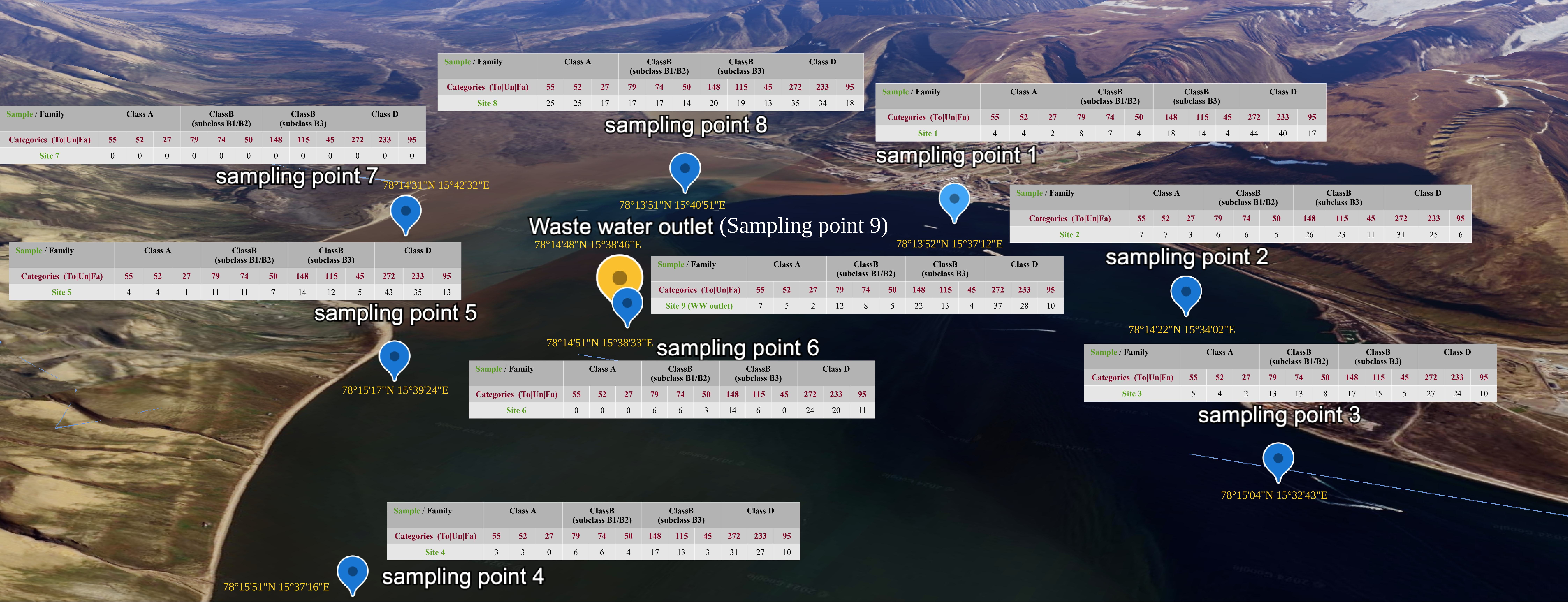


<table>
<tr><th>Sample / Family</th><th colspan="3">Class A</th><th colspan="3">ClassB (subclass B1/B2)</th><th colspan="3">ClassB (subclass B3)</th><th colspan="3">Class D</th></tr>
<tr><td>Categories (To|Un|Fa)</td><td>55</td><td>52</td><td>27</td><td>79</td><td>74</td><td>50</td><td>148</td><td>115</td><td>45</td><td>272</td><td>233</td><td>95</td></tr>
<tr><td>Site 8</td><td>25</td><td>25</td><td>17</td><td>17</td><td>17</td><td>14</td><td>20</td><td>19</td><td>13</td><td>35</td><td>34</td><td>18</td></tr>
</table>

<table>
<tr><th>Sample / Family</th><th colspan="3">Class A</th><th colspan="3">ClassB (subclass B1/B2)</th><th colspan="3">ClassB (subclass B3)</th><th colspan="3">Class D</th></tr>
<tr><td>Categories (To|Un|Fa)</td><td>55</td><td>52</td><td>27</td><td>79</td><td>74</td><td>50</td><td>148</td><td>115</td><td>45</td><td>272</td><td>233</td><td>95</td></tr>
<tr><td>Site 7</td><td>0</td><td>0</td><td>0</td><td>0</td><td>0</td><td>0</td><td>0</td><td>0</td><td>0</td><td>0</td><td>0</td><td>0</td></tr>
</table>

<table>
<tr><th>Sample / Family</th><th colspan="3">Class A</th><th colspan="3">ClassB (subclass B1/B2)</th><th colspan="3">ClassB (subclass B3)</th><th colspan="3">Class D</th></tr>
<tr><td>Categories (To|Un|Fa)</td><td>55</td><td>52</td><td>27</td><td>79</td><td>74</td><td>50</td><td>148</td><td>115</td><td>45</td><td>272</td><td>233</td><td>95</td></tr>
<tr><td>Site 1</td><td>4</td><td>4</td><td>2</td><td>8</td><td>7</td><td>4</td><td>18</td><td>14</td><td>4</td><td>44</td><td>40</td><td>17</td></tr>
</table>

<table>
<tr><th>Sample / Family</th><th colspan="3">Class A</th><th colspan="3">ClassB (subclass B1/B2)</th><th colspan="3">ClassB (subclass B3)</th><th colspan="3">Class D</th></tr>
<tr><td>Categories (To|Un|Fa)</td><td>55</td><td>52</td><td>27</td><td>79</td><td>74</td><td>50</td><td>148</td><td>115</td><td>45</td><td>272</td><td>233</td><td>95</td></tr>
<tr><td>Site 2</td><td>7</td><td>7</td><td>3</td><td>6</td><td>6</td><td>5</td><td>26</td><td>23</td><td>11</td><td>31</td><td>25</td><td>6</td></tr>
</table>

<table>
<tr><th>Sample / Family</th><th colspan="3">Class A</th><th colspan="3">ClassB (subclass B1/B2)</th><th colspan="3">ClassB (subclass B3)</th><th colspan="3">Class D</th></tr>
<tr><td>Categories (To|Un|Fa)</td><td>55</td><td>52</td><td>27</td><td>79</td><td>74</td><td>50</td><td>148</td><td>115</td><td>45</td><td>272</td><td>233</td><td>95</td></tr>
<tr><td>Site 5</td><td>4</td><td>4</td><td>1</td><td>11</td><td>11</td><td>7</td><td>14</td><td>12</td><td>5</td><td>43</td><td>35</td><td>13</td></tr>
</table>

<table>
<tr><th>Sample / Family</th><th colspan="3">Class A</th><th colspan="3">ClassB (subclass B1/B2)</th><th colspan="3">ClassB (subclass B3)</th><th colspan="3">Class D</th></tr>
<tr><td>Categories (To|Un|Fa)</td><td>55</td><td>52</td><td>27</td><td>79</td><td>74</td><td>50</td><td>148</td><td>115</td><td>45</td><td>272</td><td>233</td><td>95</td></tr>
<tr><td>Site 9 (WW outlet)</td><td>7</td><td>5</td><td>2</td><td>12</td><td>8</td><td>5</td><td>22</td><td>13</td><td>4</td><td>37</td><td>28</td><td>10</td></tr>
</table>

<table>
<tr><th>Sample / Family</th><th colspan="3">Class A</th><th colspan="3">ClassB (subclass B1/B2)</th><th colspan="3">ClassB (subclass B3)</th><th colspan="3">Class D</th></tr>
<tr><td>Categories (To|Un|Fa)</td><td>55</td><td>52</td><td>27</td><td>79</td><td>74</td><td>50</td><td>148</td><td>115</td><td>45</td><td>272</td><td>233</td><td>95</td></tr>
<tr><td>Site 3</td><td>5</td><td>4</td><td>2</td><td>13</td><td>13</td><td>8</td><td>17</td><td>15</td><td>5</td><td>27</td><td>24</td><td>10</td></tr>
</table>

<table>
<tr><th>Sample / Family</th><th colspan="3">Class A</th><th colspan="3">ClassB (subclass B1/B2)</th><th colspan="3">ClassB (subclass B3)</th><th colspan="3">Class D</th></tr>
<tr><td>Categories (To|Un|Fa)</td><td>55</td><td>52</td><td>27</td><td>79</td><td>74</td><td>50</td><td>148</td><td>115</td><td>45</td><td>272</td><td>233</td><td>95</td></tr>
<tr><td>Site 6</td><td>0</td><td>0</td><td>0</td><td>6</td><td>6</td><td>3</td><td>14</td><td>6</td><td>0</td><td>24</td><td>20</td><td>11</td></tr>
</table>

<table>
<tr><th>Sample / Family</th><th colspan="3">Class A</th><th colspan="3">ClassB (subclass B1/B2)</th><th colspan="3">ClassB (subclass B3)</th><th colspan="3">Class D</th></tr>
<tr><td>Categories (To|Un|Fa)</td><td>55</td><td>52</td><td>27</td><td>79</td><td>74</td><td>50</td><td>148</td><td>115</td><td>45</td><td>272</td><td>233</td><td>95</td></tr>
<tr><td>Site 4</td><td>3</td><td>3</td><td>0</td><td>6</td><td>6</td><td>4</td><td>17</td><td>13</td><td>3</td><td>31</td><td>27</td><td>10</td></tr>
</table>

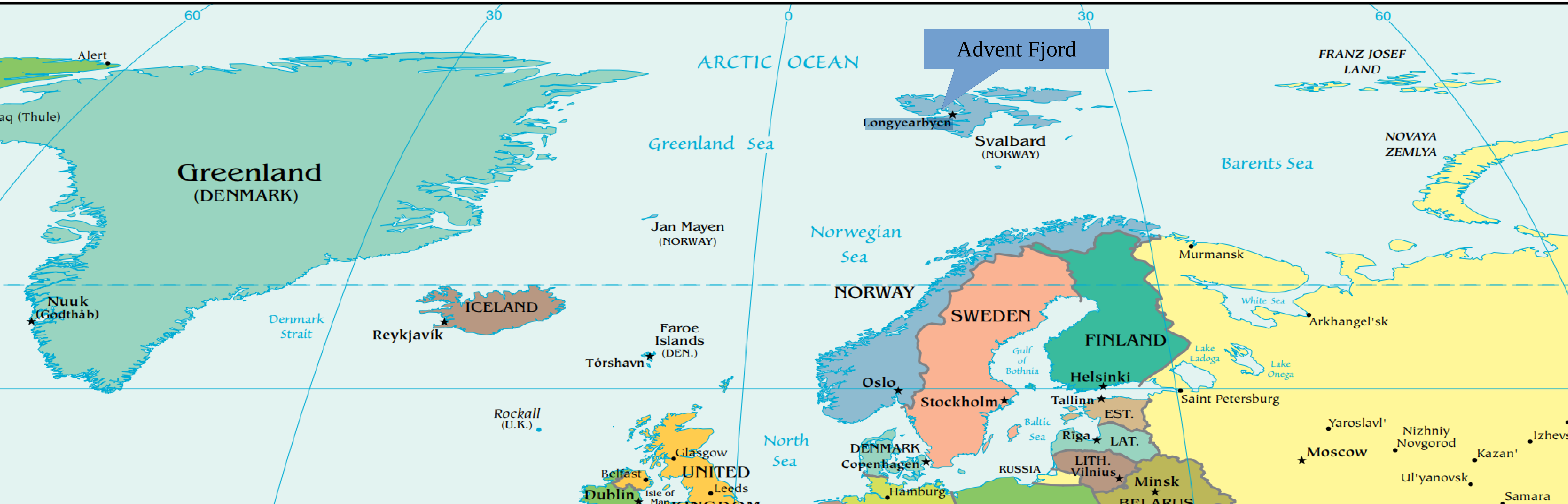
