## Supplementary 3a for "Identification of clinical carbapenemases, 478 novel β-lactamases and putative seven new bacterial phyla from wastewater contaminated high arctic fjord sediments"

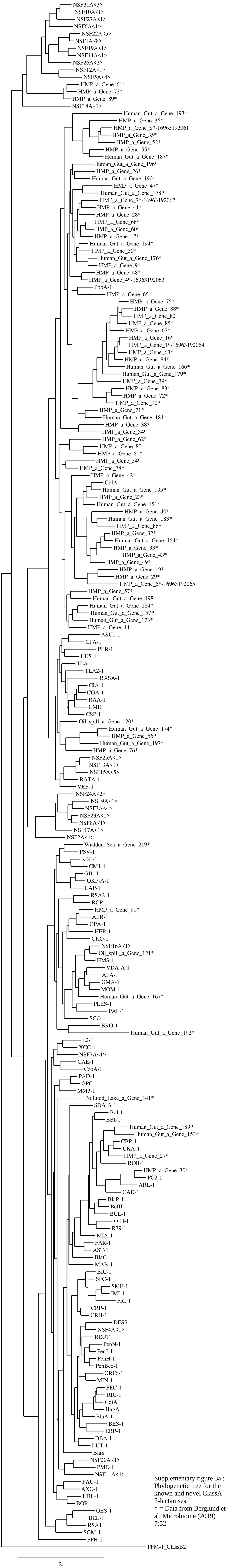


Supplementary figure 3a : Phylogenetic tree for the known and novel ClassA β-lactamses.
* = Data from Berglund et al. Microbiome (2019) 7:52
