## Supplementary 4 for "Identification of clinical carbapenemases, 478 novel β-lactamases and putative seven new bacterial phyla from wastewater contaminated high arctic fjord sediments"

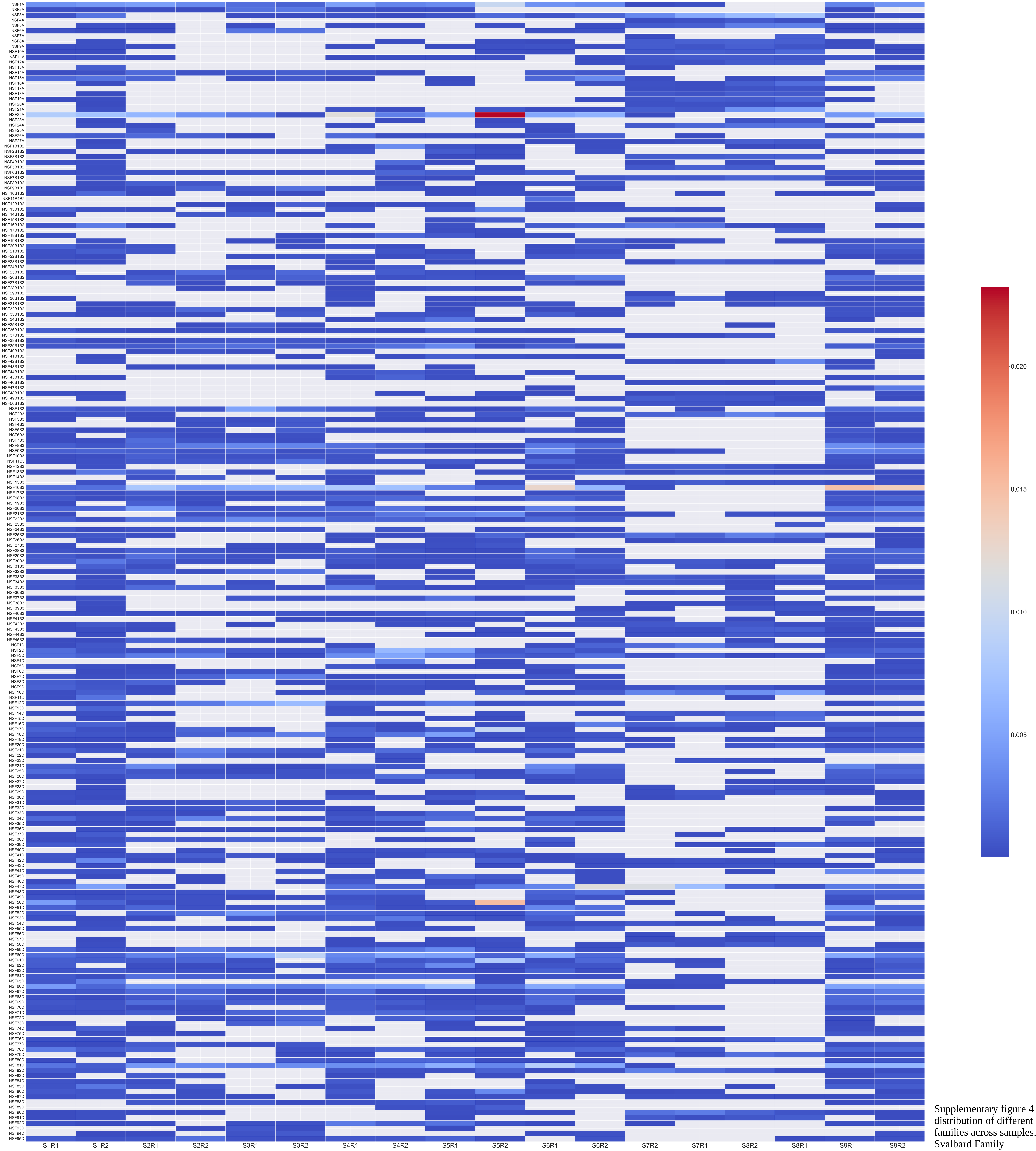


Supplementary figure 4 : Heatmap showing distribution of different novel β-lactamases families across samples. NSF = Novel Svalbard Family
