## Supplementary 5 for "Identification of clinical carbapenemases, 478 novel β-lactamases and putative seven new bacterial phyla from wastewater contaminated high arctic fjord sediments"

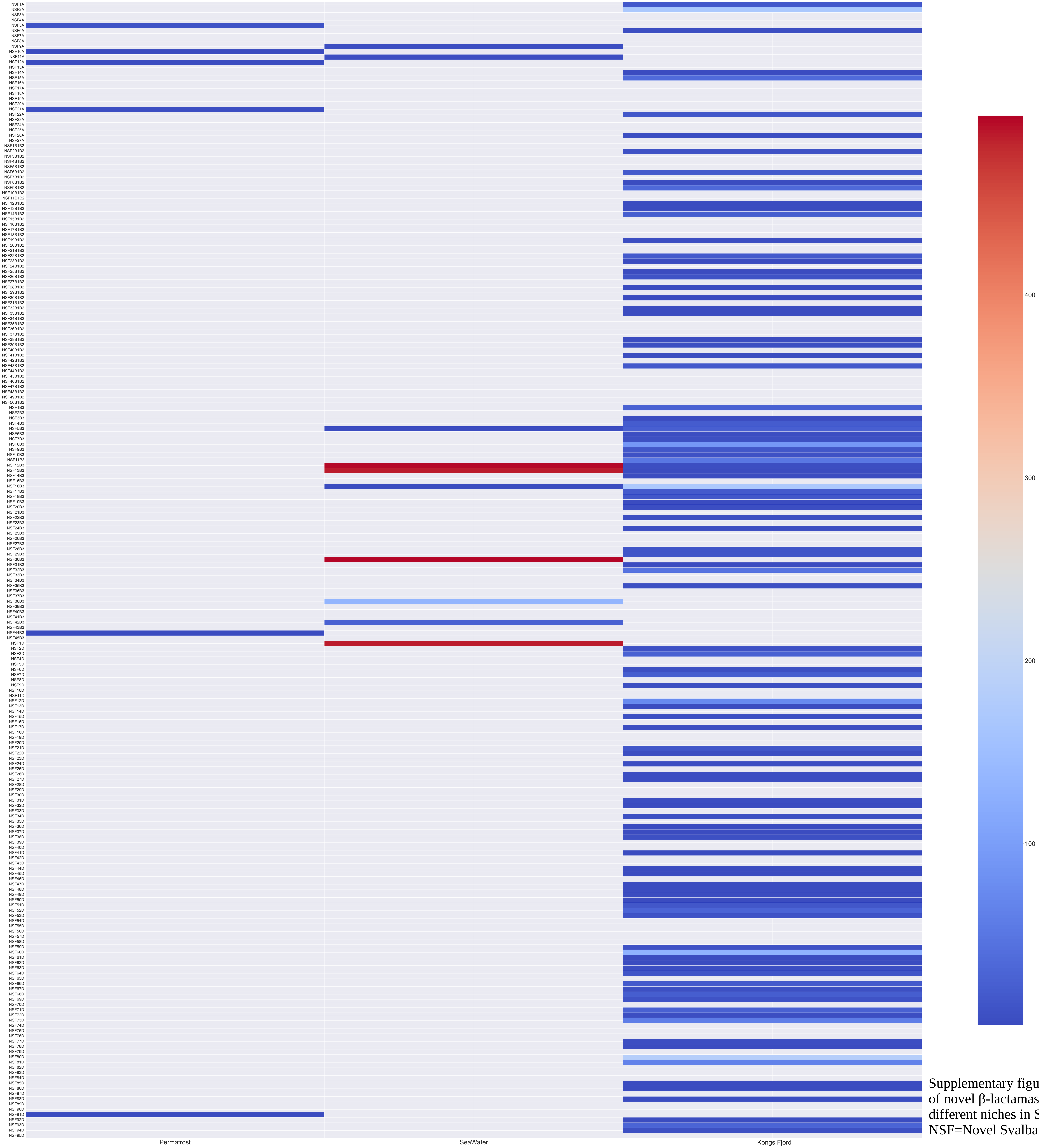


Supplementary figure 5 : Distribution of novel β-lactamase families in different niches in Svalbard. NSF=Novel Svalbard Family
