## Supplementary table2 for "Identification of clinical carbapenemases, 478 novel β-lactamases and putative seven new bacterial phyla from wastewater contaminated high arctic fjord sediments"

### Metabolic pathways detected in 644 MAGs. The results have hyperlink to the metacyc pathways for the pathways and BRENDA for enzymes used in the pathways.

| Pathway | Description | #Functions | #Annotated | Functions |
| --- | --- | --- | --- | --- |
| P101-PWY | ectoine biosynthesis | 5 | 5 | EC.1.2.1.11 EC.2.3.1.178 EC.2.6.1.76 EC.2.7.2.4 EC.4.2.1.108 |
| P105-PWY | TCA cycle IV (2-oxoglutarate decarboxylase) | 10 | 8 | EC.1.1.1.37 EC.1.1.1.42 EC.1.3.5.1 EC.2.3.3.1 EC.2.3.3.9 EC.4.1.1.71 EC.4.1.3.1 EC.4.2.1.2 |
| P108-PWY | pyruvate fermentation to propanoate I | 7 | 3 | EC.2.1.3.1 EC.5.1.99.1 EC.5.4.99.2 |
| P121-PWY | adenine and adenosine salvage I | 2 | 2 | EC.2.4.2.1 EC.2.4.2.7 |
| P122-PWY | heterolactic fermentation | 18 | 16 | EC.1.1.1.1 EC.1.1.1.27 EC.1.1.1.28 EC.1.1.1.49 EC.1.2.1.10 EC.1.2.1.12 EC.2.3.1.8 EC.2.7.1.2 EC.2.7.1.4 EC.2.7.1.40 EC.2.7.2.3 EC.3.1.1.31 EC.4.1.2.9 EC.4.2.1.11 EC.5.1.3.1 EC.5.3.1.9 |
| P124-PWY | Bifidobacterium shunt | 15 | 4 | EC.2.2.1.1 EC.2.2.1.2 EC.2.7.2.1 EC.5.3.1.6 |
| P141-PWY | atrazine degradation I (aerobic) | 3 | 3 | EC.3.5.99.3 EC.3.5.99.4 EC.3.8.1.8 |
| P142-PWY | pyruvate fermentation to acetate I | 1 | 1 | EC.1.2.7.1 |
| P162-PWY | L-glutamate degradation V (via hydroxyglutarate) | 12 | 9 | EC.1.1.1.35 EC.1.3.8.1 EC.1.4.1.2 EC.2.3.1.9 EC.2.8.3.1 EC.2.8.3.12 EC.2.8.3.8 EC.4.1.1.70 EC.6.2.1.13 |
| P163-PWY | L-lysine fermentation to acetate and butanoate | 10 | 5 | EC.1.4.1.11 EC.2.8.3.9 EC.4.3.1.14 EC.5.4.3.2 EC.5.4.3.3 |
| P164-PWY | purine nucleobases degradation I (anaerobic) | 12 | 7 | EC.1.17.1.4 EC.1.2.1.2 EC.1.21.4.2 EC.3.5.1.10 EC.3.5.4.3 EC.3.5.4.9 EC.4.3.1.4 |
| P181-PWY | nicotine degradation I (pyridine pathway) | 10 | 1 | EC.2.6.1.19 |
| P183-PWY | catechol degradation to 2-oxopent-4-enoate I | 2 | 2 | EC.1.13.11.2 EC.3.7.1.9 |
| P184-PWY | protocatechuate degradation I (meta-cleavage pathway) | 8 | 6 | EC.1.1.1.312 EC.1.13.11.8 EC.3.1.1.57 EC.4.1.1.3 EC.4.1.3.17 EC.4.2.1.83 |
| P185-PWY | formaldehyde assimilation III (dihydroxyacetone cycle) | 11 | 4 | EC.2.7.1.29 EC.3.1.3.11 EC.4.1.2.13 EC.5.3.1.1 |
| P2-PWY | citrate lyase activation | 5 | 2 | EC.2.7.7.61 EC.6.2.1.22 |
| P221-PWY | octane oxidation | 5 | 4 | EC.1.14.15.3 EC.1.18.1.1 EC.1.2.1.3 EC.6.2.1.3 |
| P224-PWY | sulfate reduction V (dissimilatory) | 3 | 3 | EC.1.8.99.2 EC.1.8.99.3 EC.2.7.7.4 |
| P23-PWY | reductive TCA cycle I | 10 | 5 | EC.1.2.7.3 EC.2.3.3.8 EC.2.7.9.2 EC.4.1.1.31 EC.6.2.1.5 |
| P241-PWY | coenzyme B biosynthesis | 7 | 2 | EC.2.3.3.14 EC.4.2.1.36 |
| P261-PWY | coenzyme M biosynthesis I | 4 | 3 | EC.3.1.3.71 EC.4.1.1.79 EC.4.4.1.19 |
| P281-PWY | 3-phenylpropanoate degradation | 1 | 1 | EC.1.14.13.58 |
| P283-PWY | hydrogen oxidation I (aerobic) | 1 | 1 | EC.1.12.99.6 |
| P3-PWY | gallate degradation III (anaerobic) | 12 | 1 | EC.1.97.1.2 |
| P302-PWY | L-sorbose degradation | 2 | 1 | EC.1.1.1.140 |
| P303-PWY | ammonia oxidation II (anaerobic) | 3 | 1 | EC.1.7.2.1 |
| P321-PWY | benzoyl-CoA degradation III (anaerobic) | 5 | 1 | EC.1.3.7.8 |
| P341-PWY | glycolysis V (Pyrococcus) | 9 | 1 | EC.1.2.7.6 |
| P345-PWY | aldoxime degradation | 3 | 2 | EC.3.5.1.19 EC.4.2.1.84 |
| P42-PWY | incomplete reductive TCA cycle | 7 | 1 | EC.6.4.1.1 |
| P483-PWY | phosphonoacetate degradation | 1 | 1 | EC.3.11.1.2 |
| P541-PWY | glycine betaine biosynthesis IV (from glycine) | 2 | 1 | EC.2.1.1.20 |
| P542-PWY | choline-O-sulfate degradation | 1 | 1 | EC.3.1.6.6 |
| P562-PWY | myo-inositol degradation I | 7 | 4 | EC.1.1.1.18 EC.2.7.1.92 EC.4.1.2.29 EC.4.2.1.44 |
| P621-PWY | nylon-6 oligomer degradation | 6 | 2 | EC.3.5.1.46 EC.3.5.2.12 |
| P641-PWY | phenylmercury acetate degradation | 2 | 2 | EC.1.16.1.1 EC.4.99.1.2 |
| PANTO-PWY | phosphopantothenate biosynthesis I | 4 | 4 | EC.1.1.1.169 EC.2.1.2.11 EC.2.7.1.33 EC.6.3.2.1 |
| PARATHION-DEGRADATION-PWY | parathion degradation | 1 | 1 | EC.3.1.8.1 |
| PCEDEG-PWY | tetrachloroethene degradation | 2 | 1 | EC.1.97.1.8 |
| PCPDEG-PWY | pentachlorophenol degradation | 2 | 1 | EC.1.14.13.50 |
| PEPTIDOGLYCANSYN-PWY | peptidoglycan biosynthesis I (meso-diaminopimelate containing) | 2 | 2 | EC.2.4.1.227 EC.2.7.8.13 |
| PHENOLDEG-PWY | phenol degradation II (anaerobic) | 2 | 2 | EC.1.3.7.9 EC.6.2.1.27 |
| PHENYLALANINE-DEG1-PWY | L-phenylalanine degradation I (aerobic) | 3 | 2 | EC.1.14.16.1 EC.4.2.1.96 |
| PHESYN | L-phenylalanine biosynthesis I | 3 | 3 | EC.2.6.1.57 EC.4.2.1.51 EC.5.4.99.5 |
| PHOSLIPSYN2-PWY | superpathway of phospholipid biosynthesis II (plants) | 3 | 2 | EC.2.7.8.11 EC.2.7.8.8 |
| PHOSPHONOTASE-PWY | 2-aminoethylphosphonate degradation I | 3 | 2 | EC.2.6.1.37 EC.3.11.1.1 |
| PLPSAL-PWY | pyridoxal 5'-phosphate salvage I | 2 | 2 | EC.1.4.3.5 EC.2.7.1.35 |
| POLYAMINSYN3-PWY | superpathway of polyamine biosynthesis II | 1 | 1 | EC.2.5.1.22 |
| PPGPPMET-PWY | ppGpp biosynthesis | 4 | 4 | EC.2.7.4.6 EC.2.7.6.5 EC.3.1.7.2 EC.3.6.1.40 |
| PROPIONMET-PWY | propanoyl CoA degradation I | 3 | 1 | EC.6.4.1.3 |
| PROSYN-PWY | L-proline biosynthesis I | 3 | 3 | EC.1.2.1.41 EC.1.5.1.2 EC.2.7.2.11 |
| PROTOCATECHUATE-ORTHO-CLEAVAGE-PWY | protocatechuate degradation II (ortho-cleavage pathway) | 4 | 4 | EC.1.13.11.3 EC.3.1.1.24 EC.4.1.1.44 EC.5.5.1.2 |
| PUTDEG-PWY | putrescine degradation I | 1 | 1 | EC.1.2.1.19 |
| PWY-0 | putrescine degradation III | 4 | 1 | EC.2.3.1.57 |
| PWY-1001 | cellulose biosynthesis | 1 | 1 | EC.2.4.1.12 |
| PWY-101 | photosynthesis light reactions | 4 | 2 | EC.1.10.9.1 EC.1.18.1.2 |
| PWY-1042 | glycolysis IV (plant cytosol) | 10 | 3 | EC.1.2.1.9 EC.2.7.1.11 EC.2.7.1.90 |
| PWY-1081 | homogalacturonan degradation | 2 | 2 | EC.3.1.1.11 EC.3.2.1.15 |
| PWY-1121 | suberin monomers biosynthesis | 14 | 3 | EC.2.1.1.104 EC.3.1.2.2 EC.6.2.1.12 |
| PWY-1263 | taurine degradation I | 1 | 1 | EC.2.6.1.77 |
| PWY-1264 | taurine degradation II | 1 | 1 | EC.1.4.99.2 |
| PWY-1269 | CMP-3-deoxy-D-manno-octulosonate biosynthesis I | 4 | 4 | EC.2.5.1.55 EC.2.7.7.38 EC.3.1.3.45 EC.5.3.1.13 |
| PWY-1281 | sulfoacetaldehyde degradation I | 2 | 1 | EC.2.3.3.15 |
| PWY-1341 | phenylacetate degradation II (anaerobic) | 4 | 2 | EC.1.17.5.1 EC.6.2.1.30 |
| PWY-1361 | benzoyl-CoA degradation I (aerobic) | 6 | 1 | EC.2.3.1.174 |
| PWY-1422 | vitamin E biosynthesis (tocopherols) | 5 | 1 | EC.1.13.11.27 |
| PWY-1501 | mandelate degradation I | 5 | 3 | EC.1.1.99.31 EC.4.1.1.7 EC.5.1.2.2 |
| PWY-1622 | formaldehyde assimilation I (serine pathway) | 10 | 4 | EC.1.1.1.81 EC.2.1.2.1 EC.2.6.1.45 EC.4.1.3.24 |
| PWY-1641 | methane oxidation to methanol I | 1 | 1 | EC.1.14.13.25 |
| PWY-1722 | formate reduction to 5,10-methylenetetrahydrofolate | 3 | 2 | EC.1.5.1.5 EC.6.3.4.3 |
| PWY-1723 | formaldehyde oxidation V (H4MPT pathway) | 3 | 1 | EC.3.5.4.27 |
| PWY-1781 | β-alanine degradation II | 2 | 1 | EC.2.6.1.18 |
| PWY-1801 | formaldehyde oxidation II (glutathione-dependent) | 3 | 3 | EC.1.1.1.284 EC.3.1.2.12 EC.4.4.1.22 |
| PWY-181 | photorespiration | 8 | 4 | EC.1.1.1.29 EC.1.1.3.15 EC.2.7.1.31 EC.3.1.3.18 |
| PWY-2 | putrescine degradation IV | 3 | 1 | EC.2.6.1.82 |
| PWY-2002 | isoflavonoid biosynthesis I | 5 | 1 | EC.5.5.1.6 |
| PWY-2161 | folate polyglutamylation | 3 | 1 | EC.6.3.2.17 |
| PWY-2201 | folate transformations I | 11 | 3 | EC.1.5.1.20 EC.2.1.1.13 EC.6.3.3.2 |
| PWY-2221 | Entner-Doudoroff pathway III (semi-phosphorylative) | 13 | 3 | EC.2.7.1.45 EC.3.1.1.17 EC.4.1.2.14 |
| PWY-2301 | myo-inositol biosynthesis | 2 | 2 | EC.3.1.3.25 EC.5.5.1.4 |
| PWY-2361 | 3-oxoadipate degradation | 2 | 1 | EC.2.8.3.6 |
| PWY-241 | C4 photosynthetic carbon assimilation cycle, NADP-ME type | 5 | 3 | EC.1.1.1.40 EC.2.7.9.1 EC.4.2.1.1 |
| PWY-2421 | indole-3-acetate degradation VIII (bacterial) | 4 | 1 | EC.1.14.12.1 |
| PWY-2503 | benzoate degradation I (aerobic) | 2 | 2 | EC.1.14.12.10 EC.1.3.1.25 |
| PWY-2541 | plant sterol biosynthesis | 14 | 4 | EC.1.14.13.70 EC.1.3.1.21 EC.1.3.1.70 EC.2.1.1.41 |
| PWY-2582 | brassinosteroid biosynthesis II | 10 | 1 | EC.5.3.3.1 |
| PWY-2622 | trehalose biosynthesis IV | 1 | 1 | EC.5.4.99.16 |
| PWY-2661 | trehalose biosynthesis V | 3 | 2 | EC.3.2.1.141 EC.5.4.99.15 |
| PWY-2721 | trehalose degradation III | 2 | 1 | EC.5.4.2.6 |
| PWY-2722 | trehalose degradation IV | 4 | 2 | EC.2.4.1.64 EC.2.7.1.1 |
| PWY-2723 | trehalose degradation V | 6 | 3 | EC.5.1.3.15 EC.5.1.3.3 EC.5.4.2.2 |
| PWY-2781 | cis-zeatin biosynthesis | 1 | 1 | EC.2.5.1.75 |
| PWY-282 | cuticular wax biosynthesis | 5 | 2 | EC.1.2.1.50 EC.4.1.99.5 |
| PWY-283 | benzoate degradation II (aerobic and anaerobic) | 1 | 1 | EC.6.2.1.25 |
| PWY-2941 | L-lysine biosynthesis II | 8 | 6 | EC.1.17.1.8 EC.2.3.1.89 EC.3.5.1.47 EC.4.1.1.20 EC.4.3.3.7 EC.5.1.1.7 |
| PWY-3081 | L-lysine biosynthesis V | 9 | 1 | EC.2.6.1.39 |
| PWY-31 | canavanine degradation | 2 | 1 | EC.3.5.3.1 |
| PWY-3121 | linamarin degradation | 2 | 1 | EC.3.2.1.21 |
| PWY-3161 | indole-3-acetate biosynthesis III (bacteria) | 2 | 2 | EC.1.13.12.3 EC.3.5.1.4 |
| PWY-3162 | L-tryptophan degradation V (side chain pathway) | 6 | 1 | EC.1.2.1.5 |
| PWY-3181 | L-tryptophan degradation VI (via tryptamine) | 3 | 1 | EC.4.1.1.28 |
| PWY-3221 | dTDP-L-rhamnose biosynthesis II | 2 | 2 | EC.2.7.7.24 EC.4.2.1.46 |
| PWY-3261 | UDP-L-rhamnose biosynthesis | 1 | 1 | EC.4.2.1.76 |
| PWY-3341 | L-proline biosynthesis III | 2 | 1 | EC.2.6.1.13 |
| PWY-3385 | choline biosynthesis I | 4 | 1 | EC.2.1.1.103 |
| PWY-3461 | L-tyrosine biosynthesis II | 3 | 1 | EC.1.3.1.78 |
| PWY-3462 | L-phenylalanine biosynthesis II | 3 | 1 | EC.4.2.1.91 |
| PWY-3561 | choline biosynthesis III | 3 | 3 | EC.2.7.7.15 EC.2.7.8.2 EC.3.1.4.4 |
| PWY-3581 | (S)-reticuline biosynthesis I | 10 | 1 | EC.4.1.1.25 |
| PWY-3602 | L-carnitine degradation II | 1 | 1 | EC.1.1.1.108 |
| PWY-3621 | γ-butyrobetaine degradation | 1 | 1 | EC.1.14.11.1 |
| PWY-3641 | L-carnitine degradation III | 4 | 1 | EC.1.1.1.39 |
| PWY-3661 | glycine betaine degradation I | 5 | 3 | EC.1.5.3.1 EC.1.5.8.4 EC.2.1.1.5 |
| PWY-3722 | glycine betaine biosynthesis II (Gram-positive bacteria) | 2 | 1 | EC.1.2.1.8 |
| PWY-3781 | aerobic respiration I (cytochrome c) | 4 | 3 | EC.1.10.2.2 EC.1.6.5.3 EC.1.9.3.1 |
| PWY-3801 | sucrose degradation II (sucrose synthase) | 7 | 2 | EC.2.4.1.13 EC.2.7.7.9 |
| PWY-381 | nitrate reduction II (assimilatory) | 3 | 3 | EC.1.7.1.1 EC.1.7.7.1 EC.6.3.1.2 |
| PWY-3821 | galactose degradation III | 4 | 3 | EC.2.7.1.6 EC.2.7.7.10 EC.5.1.3.2 |
| PWY-3841 | folate transformations II | 9 | 2 | EC.1.5.1.3 EC.2.1.1.45 |
| PWY-3861 | mannitol degradation II | 3 | 2 | EC.1.1.1.255 EC.5.3.1.8 |
| PWY-3941 | β-alanine biosynthesis II | 6 | 2 | EC.3.1.2.4 EC.6.2.1.17 |
| PWY-3982 | uracil degradation I (reductive) | 3 | 3 | EC.1.3.1.2 EC.3.5.1.6 EC.3.5.2.2 |
| PWY-40 | putrescine biosynthesis I | 2 | 2 | EC.3.5.3.11 EC.4.1.1.19 |
| PWY-401 | galactolipid biosynthesis I | 3 | 1 | EC.2.4.1.46 |
| PWY-4041 | γ-glutamyl cycle | 4 | 3 | EC.2.3.2.2 EC.2.3.2.4 EC.3.5.2.9 |
| PWY-4061 | glutathione-mediated detoxification I | 5 | 1 | EC.2.5.1.18 |
| PWY-4081 | glutathione redox reactions I | 3 | 2 | EC.1.11.1.9 EC.1.8.1.7 |
| PWY-4101 | D-sorbitol degradation I | 2 | 1 | EC.1.1.1.14 |
| PWY-4121 | glutathionylspermidine biosynthesis | 2 | 1 | EC.6.3.1.8 |
| PWY-4202 | arsenate detoxification I (glutaredoxin) | 4 | 2 | EC.1.20.4.1 EC.2.1.1.137 |
| PWY-4261 | glycerol degradation I | 2 | 2 | EC.1.1.5.3 EC.2.7.1.30 |
| PWY-43 | putrescine biosynthesis II | 3 | 2 | EC.3.5.1.53 EC.3.5.3.12 |
| PWY-4321 | L-glutamate degradation IV | 5 | 2 | EC.1.2.1.24 EC.4.1.1.15 |
| PWY-4341 | L-glutamate biosynthesis V | 1 | 1 | EC.1.4.7.1 |
| PWY-4361 | S-methyl-5-thio-α-D-ribose 1-phosphate degradation | 7 | 3 | EC.1.13.11.54 EC.4.2.1.109 EC.5.3.1.23 |
| PWY-4381 | fatty acid biosynthesis initiation I | 5 | 5 | EC.2.3.1.180 EC.2.3.1.39 EC.2.3.1.85 EC.2.3.1.86 EC.6.4.1.2 |
| PWY-46 | putrescine biosynthesis III | 1 | 1 | EC.4.1.1.17 |
| PWY-4621 | arsenate detoxification II (glutaredoxin) | 2 | 1 | EC.3.6.3.16 |
| PWY-4702 | phytate degradation I | 4 | 2 | EC.3.1.3.26 EC.3.1.3.8 |
| PWY-4722 | creatinine degradation II | 5 | 3 | EC.1.5.8.3 EC.3.5.1.59 EC.3.5.2.14 |
| PWY-4821 | UDP-D-xylose biosynthesis | 1 | 1 | EC.4.1.1.35 |
| PWY-4841 | UDP-α-D-glucuronate biosynthesis (from myo-inositol) | 3 | 1 | EC.1.13.99.1 |
| PWY-4861 | UDP-D-galacturonate biosynthesis I (from UDP-D-glucuronate) | 1 | 1 | EC.5.1.3.6 |
| PWY-4921 | protein citrullination | 1 | 1 | EC.3.5.3.15 |
| PWY-4942 | sterculate biosynthesis | 2 | 1 | EC.2.1.1.79 |
| PWY-4981 | L-proline biosynthesis II (from arginine) | 5 | 2 | EC.2.1.3.3 EC.3.5.3.6 |
| PWY-4983 | L-citrulline-nitric oxide cycle | 3 | 3 | EC.1.14.13.39 EC.4.3.2.1 EC.6.3.4.5 |
| PWY-4984 | urea cycle | 5 | 1 | EC.6.3.4.16 |
| PWY-5022 | 4-aminobutanoate degradation V | 8 | 2 | EC.1.1.1.61 EC.4.2.1.120 |
| PWY-5026 | indole-3-acetate biosynthesis V (bacteria and fungi) | 1 | 1 | EC.3.5.5.1 |
| PWY-5028 | L-histidine degradation II | 5 | 5 | EC.3.5.1.68 EC.3.5.2.7 EC.3.5.3.13 EC.4.2.1.49 EC.4.3.1.3 |
| PWY-5030 | L-histidine degradation III | 6 | 1 | EC.2.1.2.5 |
| PWY-5033 | nicotinate degradation II | 4 | 1 | EC.5.2.1.1 |
| PWY-5041 | S-adenosyl-L-methionine cycle II | 4 | 3 | EC.2.1.1.14 EC.2.5.1.6 EC.3.3.1.1 |
| PWY-5046 | 2-oxoisovalerate decarboxylation to isobutanoyl-CoA | 3 | 3 | EC.1.2.4.4 EC.1.8.1.4 EC.2.3.1.168 |
| PWY-5049 | rosmarinic acid biosynthesis II | 4 | 1 | EC.1.14.16.2 |
| PWY-5055 | nicotinate degradation III | 8 | 5 | EC.1.1.1.291 EC.1.17.1.5 EC.1.3.7.1 EC.4.1.3.32 EC.5.3.3.6 |
| PWY-5057 | L-valine degradation II | 3 | 1 | EC.2.6.1.42 |
| PWY-5067 | glycogen biosynthesis II (from UDP-D-Glucose) | 3 | 2 | EC.2.4.1.11 EC.2.4.1.18 |
| PWY-5068 | chlorophyll cycle | 4 | 1 | EC.2.5.1.62 |
| PWY-5074 | mevalonate degradation | 2 | 2 | EC.1.1.1.88 EC.4.1.3.4 |
| PWY-5076 | L-leucine degradation III | 4 | 1 | EC.4.1.1.1 |
| PWY-5081 | L-tryptophan degradation VIII (to tryptophol) | 4 | 1 | EC.4.1.1.74 |
| PWY-5083 | NAD/NADH phosphorylation and dephosphorylation | 7 | 3 | EC.1.6.1.1 EC.1.6.1.2 EC.2.7.1.23 |
| PWY-5084 | 2-oxoglutarate decarboxylation to succinyl-CoA | 3 | 2 | EC.1.2.4.2 EC.2.3.1.61 |
| PWY-5087 | L-glutamate degradation VI (to pyruvate) | 4 | 2 | EC.4.3.1.2 EC.5.4.99.1 |
| PWY-5097 | L-lysine biosynthesis VI | 7 | 1 | EC.2.6.1.83 |
| PWY-5098 | chlorophyll a degradation I | 4 | 1 | EC.3.1.1.14 |
| PWY-5101 | L-isoleucine biosynthesis II | 7 | 3 | EC.2.2.1.6 EC.4.2.1.35 EC.4.2.1.9 |
| PWY-5103 | L-isoleucine biosynthesis III | 5 | 1 | EC.1.1.1.86 |
| PWY-5109 | 2-methylbutanoate biosynthesis | 5 | 2 | EC.1.1.1.178 EC.4.2.1.17 |
| PWY-5110 | trigonelline biosynthesis | 1 | 1 | EC.2.1.1.7 |
| PWY-5120 | geranylgeranyl diphosphate biosynthesis | 1 | 1 | EC.2.5.1.29 |
| PWY-5122 | geranyl diphosphate biosynthesis | 1 | 1 | EC.2.5.1.1 |
| PWY-5123 | trans, trans-farnesyl diphosphate biosynthesis | 3 | 2 | EC.2.5.1.10 EC.5.3.3.2 |
| PWY-5129 | sphingolipid biosynthesis (plants) | 12 | 1 | EC.2.3.1.50 |
| PWY-5135 | xanthohumol biosynthesis | 4 | 1 | EC.2.3.1.74 |
| PWY-5136 | fatty acid β-oxidation II (peroxisome) | 5 | 2 | EC.1.3.3.6 EC.2.3.1.16 |
| PWY-5137 | fatty acid β-oxidation III (unsaturated, odd number) | 1 | 1 | EC.5.3.3.8 |
| PWY-5138 | unsaturated, even numbered fatty acid β-oxidation | 5 | 2 | EC.1.3.1.34 EC.5.1.2.3 |
| PWY-5142 | acyl-ACP thioesterase pathway | 1 | 1 | EC.3.1.2.14 |
| PWY-5147 | oleate biosynthesis I (plants) | 3 | 1 | EC.1.14.19.2 |
| PWY-5152 | leucodelphinidin biosynthesis | 3 | 1 | EC.1.1.1.219 |
| PWY-5154 | L-arginine biosynthesis III (via N-acetyl-L-citrulline) | 9 | 7 | EC.1.2.1.38 EC.2.1.3.9 EC.2.3.1.1 EC.2.6.1.11 EC.2.7.2.8 EC.3.5.1.16 EC.6.3.5.5 |
| PWY-5155 | β-alanine biosynthesis III | 1 | 1 | EC.4.1.1.11 |
| PWY-5159 | trans-4-hydroxy-L-proline degradation II | 4 | 2 | EC.1.2.1.26 EC.5.1.1.8 |
| PWY-5162 | 2-oxopentenoate degradation | 3 | 2 | EC.4.1.3.39 EC.4.2.1.80 |
| PWY-5168 | ferulate and sinapate biosynthesis | 4 | 1 | EC.1.2.1.68 |
| PWY-5169 | cyanurate degradation | 3 | 1 | EC.3.5.1.54 |
| PWY-5188 | tetrapyrrole biosynthesis I (from glutamate) | 6 | 6 | EC.1.2.1.70 EC.2.5.1.61 EC.4.2.1.24 EC.4.2.1.75 EC.5.4.3.8 EC.6.1.1.17 |
| PWY-5189 | tetrapyrrole biosynthesis II (from glycine) | 4 | 1 | EC.2.3.1.37 |
| PWY-5194 | siroheme biosynthesis | 2 | 2 | EC.1.3.1.76 EC.4.99.1.4 |
| PWY-5197 | lactate biosynthesis (archaea) | 3 | 2 | EC.1.2.1.22 EC.4.1.2.17 |
| PWY-5198 | factor 420 biosynthesis | 4 | 1 | EC.2.5.1.77 |
| PWY-5199 | factor 420 polyglutamylation | 2 | 2 | EC.6.3.2.31 EC.6.3.2.34 |
| PWY-5207 | coenzyme B/coenzyme M regeneration | 2 | 2 | EC.1.12.98.3 EC.1.8.98.1 |
| PWY-5209 | methyl-coenzyme M oxidation to CO2 | 5 | 1 | EC.1.2.99.5 |
| PWY-5265 | peptidoglycan biosynthesis II (staphylococci) | 4 | 2 | EC.2.4.1.129 EC.3.4.16.4 |
| PWY-5269 | cardiolipin biosynthesis II | 3 | 2 | EC.2.7.8.5 EC.3.1.3.27 |
| PWY-5276 | sulfite oxidation I (sulfite oxidoreductase) | 1 | 1 | EC.1.8.2.1 |
| PWY-5283 | L-lysine degradation V | 7 | 1 | EC.1.4.3.3 |
| PWY-5290 | secologanin and strictosidine biosynthesis | 11 | 1 | EC.4.3.3.2 |
| PWY-5298 | L-lysine degradation VI | 2 | 1 | EC.2.6.1.36 |
| PWY-5301 | ajmaline and sarpagine biosynthesis | 13 | 1 | EC.1.5.1.32 |
| PWY-5316 | nicotine biosynthesis | 5 | 2 | EC.1.4.3.16 EC.2.4.2.19 |
| PWY-5317 | hyoscyamine and scopolamine biosynthesis | 4 | 1 | EC.1.1.1.206 |
| PWY-5326 | sulfite oxidation IV | 1 | 1 | EC.1.8.3.1 |
| PWY-5329 | L-cysteine degradation III | 2 | 1 | EC.2.8.1.2 |
| PWY-5331 | taurine biosynthesis | 4 | 1 | EC.1.13.11.20 |
| PWY-5337 | stachyose biosynthesis | 3 | 1 | EC.2.4.1.82 |
| PWY-5340 | sulfate activation for sulfonation | 2 | 1 | EC.2.7.1.25 |
| PWY-5344 | L-homocysteine biosynthesis | 2 | 2 | EC.2.3.1.31 EC.2.5.1.49 |
| PWY-5350 | thiosulfate disproportionation III (rhodanese) | 1 | 1 | EC.2.8.1.1 |
| PWY-5367 | petroselinate biosynthesis | 6 | 2 | EC.1.1.1.100 EC.2.3.1.179 |
| PWY-5372 | carbon tetrachloride degradation II | 2 | 1 | EC.1.2.99.2 |
| PWY-5381 | pyridine nucleotide cycling (plants) | 10 | 4 | EC.2.7.7.18 EC.3.1.3.5 EC.3.6.1.22 EC.6.3.5.1 |
| PWY-5382 | hydrogen oxidation II (aerobic, NAD) | 1 | 1 | EC.1.12.1.2 |
| PWY-5384 | sucrose degradation IV (sucrose phosphorylase) | 6 | 1 | EC.2.4.1.7 |
| PWY-5386 | methylglyoxal degradation I | 3 | 2 | EC.3.1.2.6 EC.4.4.1.5 |
| PWY-5389 | 3-methylthiopropanoate biosynthesis | 1 | 1 | EC.1.13.11.53 |
| PWY-5392 | reductive TCA cycle II | 10 | 2 | EC.4.1.3.34 EC.6.4.1.7 |
| PWY-5418 | phenol degradation I (aerobic) | 1 | 1 | EC.1.14.13.7 |
| PWY-5419 | catechol degradation to 2-oxopent-4-enoate II | 4 | 1 | EC.4.1.1.77 |
| PWY-5426 | betaxanthin biosynthesis | 2 | 1 | EC.2.1.1.6 |
| PWY-5427 | naphthalene degradation (aerobic) | 6 | 2 | EC.1.14.12.12 EC.5.99.1.4 |
| PWY-5428 | m-xylene degradation to m-toluate | 1 | 1 | EC.1.1.1.90 |
| PWY-5436 | L-threonine degradation IV | 3 | 1 | EC.4.1.2.5 |
| PWY-5443 | aminopropanol phosphate biosynthesis I | 2 | 1 | EC.4.1.1.81 |
| PWY-5451 | acetone degradation I (to methylglyoxal) | 4 | 2 | EC.1.14.14.1 EC.4.1.1.4 |
| PWY-5458 | methylglyoxal degradation V | 3 | 1 | EC.1.1.2.3 |
| PWY-5461 | betanidin degradation | 1 | 1 | EC.1.11.1.7 |
| PWY-5468 | lupanine biosynthesis | 2 | 1 | EC.4.1.1.18 |
| PWY-5469 | sesamin biosynthesis | 2 | 1 | EC.1.2.1.11 |
| PWY-5480 | pyruvate fermentation to ethanol I | 3 | 1 | EC.2.3.1.54 |
| PWY-5491 | diethylphosphate degradation | 2 | 1 | EC.3.1.3.1 |
| PWY-5499 | vitamin B6 degradation | 7 | 1 | EC.1.1.1.107 |
| PWY-5506 | methanol oxidation to formaldehyde IV | 3 | 1 | EC.1.11.1.6 |
| PWY-5508 | adenosylcobalamin biosynthesis from cobyrinate a,c-diamide II | 9 | 6 | EC.1.16.8.1 EC.2.4.2.21 EC.2.5.1.17 EC.2.7.7.62 EC.2.7.8.26 EC.3.1.3.73 |
| PWY-5514 | UDP-N-acetyl-D-galactosamine biosynthesis II | 7 | 3 | EC.2.3.1.4 EC.2.7.7.23 EC.3.5.99.6 |
| PWY-5515 | L-arabinose degradation II | 3 | 2 | EC.1.1.1.12 EC.1.1.1.21 |
| PWY-5517 | L-arabinose degradation III | 5 | 3 | EC.3.1.1.15 EC.4.2.1.25 EC.4.2.1.43 |
| PWY-5521 | L-ascorbate biosynthesis III | 2 | 1 | EC.1.1.99.21 |
| PWY-5523 | 5,6-dimethylbenzimidazole biosynthesis | 3 | 2 | EC.1.5.1.39 EC.2.7.1.26 |
| PWY-5530 | sorbitol biosynthesis II | 3 | 2 | EC.1.1.99.28 EC.2.7.1.12 |
| PWY-5531 | chlorophyllide a biosynthesis II (anaerobic) | 7 | 6 | EC.1.3.1.75 EC.1.3.3.3 EC.1.3.3.4 EC.2.1.1.11 EC.4.1.1.37 EC.6.6.1.1 |
| PWY-5532 | adenosine nucleotides degradation IV | 4 | 2 | EC.2.7.4.23 EC.4.1.1.39 |
| PWY-5533 | acetone degradation II (to acetoacetate) | 2 | 1 | EC.6.4.1.6 |
| PWY-5534 | propylene degradation | 5 | 1 | EC.1.1.1.269 |
| PWY-561 | superpathway of glyoxylate cycle and fatty acid degradation | 4 | 1 | EC.4.1.1.49 |
| PWY-5629 | isopenicillin N biosynthesis | 2 | 2 | EC.1.21.3.1 EC.6.3.2.26 |
| PWY-5630 | penicillin K biosynthesis | 1 | 1 | EC.2.3.1.164 |
| PWY-5631 | deacetylcephalosporin C biosynthesis | 3 | 1 | EC.5.1.1.17 |
| PWY-5633 | cephamycin C biosynthesis | 1 | 1 | EC.2.1.3.7 |
| PWY-5642 | 2,4-dinitrotoluene degradation | 3 | 1 | EC.1.2.1.27 |
| PWY-5647 | 2-nitrobenzoate degradation I | 5 | 4 | EC.1.13.11.6 EC.1.2.1.32 EC.3.5.99.5 EC.4.1.1.45 |
| PWY-5651 | L-tryptophan degradation to 2-amino-3-carboxymuconate semialdehyde | 6 | 4 | EC.1.13.11.11 EC.1.14.13.9 EC.3.5.1.9 EC.3.7.1.3 |
| PWY-5656 | mannosylglycerate biosynthesis I | 2 | 1 | EC.3.1.3.70 |
| PWY-5659 | GDP-mannose biosynthesis | 4 | 2 | EC.2.7.7.13 EC.5.4.2.8 |
| PWY-5663 | tetrahydrobiopterin biosynthesis I | 3 | 3 | EC.1.1.1.153 EC.3.5.4.16 EC.4.2.3.12 |
| PWY-5667 | CDP-diacylglycerol biosynthesis I | 4 | 4 | EC.1.1.1.94 EC.2.3.1.15 EC.2.3.1.51 EC.2.7.7.41 |
| PWY-5669 | phosphatidylethanolamine biosynthesis I | 2 | 1 | EC.4.1.1.65 |
| PWY-5670 | epoxysqualene biosynthesis | 2 | 2 | EC.1.14.13.132 EC.2.5.1.21 |
| PWY-5674 | nitrate reduction IV (dissimilatory) | 2 | 1 | EC.1.7.2.2 |
| PWY-5675 | nitrate reduction V (assimilatory) | 4 | 2 | EC.1.4.1.4 EC.1.7.1.4 |
| PWY-5676 | acetyl-CoA fermentation to butanoate II | 6 | 2 | EC.1.1.1.36 EC.4.2.1.55 |
| PWY-5679 | clavulanate biosynthesis | 4 | 1 | EC.3.5.3.22 |
| PWY-5686 | UMP biosynthesis | 6 | 5 | EC.1.3.5.2 EC.2.1.3.2 EC.2.4.2.10 EC.3.5.2.3 EC.4.1.1.23 |
| PWY-5690 | TCA cycle II (plants and fungi) | 7 | 1 | EC.1.1.1.41 |
| PWY-5691 | urate degradation to allantoin I | 3 | 2 | EC.1.7.3.3 EC.3.5.2.17 |
| PWY-5694 | allantoin degradation to glyoxylate I | 1 | 1 | EC.4.3.2.3 |
| PWY-5695 | urate biosynthesis/inosine 5'-phosphate degradation | 4 | 1 | EC.1.1.1.205 |
| PWY-5697 | allantoin degradation to ureidoglycolate I (urea producing) | 2 | 2 | EC.3.5.2.5 EC.3.5.3.4 |
| PWY-5703 | urea degradation I | 2 | 1 | EC.6.3.4.6 |
| PWY-5704 | urea degradation II | 1 | 1 | EC.3.5.1.5 |
| PWY-5723 | Rubisco shunt | 9 | 1 | EC.2.7.1.19 |
| PWY-5738 | GDP-6-deoxy-D-talose biosynthesis | 3 | 1 | EC.4.2.1.47 |
| PWY-5742 | L-arginine degradation IX (arginine:pyruvate transaminase pathway) | 4 | 2 | EC.3.5.3.7 EC.4.1.1.75 |
| PWY-5743 | 3-hydroxypropanoate cycle | 11 | 2 | EC.1.1.1.298 EC.1.3.1.84 |
| PWY-5747 | 2-methylcitrate cycle II | 5 | 3 | EC.2.3.3.5 EC.4.1.3.30 EC.4.2.1.99 |
| PWY-5751 | phenylethanol biosynthesis | 5 | 1 | EC.1.4.3.21 |
| PWY-5754 | 4-hydroxybenzoate biosynthesis I (eukaryotes) | 5 | 1 | EC.3.1.2.23 |
| PWY-5755 | 4-hydroxybenzoate biosynthesis II (microbes) | 1 | 1 | EC.4.1.3.40 |
| PWY-5757 | fosfomycin biosynthesis | 5 | 2 | EC.4.1.1.82 EC.5.4.2.9 |
| PWY-5766 | L-glutamate degradation X | 1 | 1 | EC.1.4.1.3 |
| PWY-5768 | pyruvate fermentation to acetate VIII | 2 | 1 | EC.1.2.1.4 |
| PWY-5785 | di-trans,poly-cis-undecaprenyl phosphate biosynthesis | 1 | 1 | EC.2.5.1.31 |
| PWY-5791 | 1,4-dihydroxy-2-naphthoate biosynthesis II (plants) | 7 | 5 | EC.2.2.1.9 EC.4.1.3.36 EC.4.2.99.20 EC.5.4.4.2 EC.6.2.1.26 |
| PWY-5807 | heptaprenyl diphosphate biosynthesis | 1 | 1 | EC.2.5.1.30 |
| PWY-5833 | CDP-4-dehydro-3,6-dideoxy-D-glucose biosynthesis | 3 | 2 | EC.2.7.7.33 EC.4.2.1.45 |
| PWY-5834 | CDP-tyvelose biosynthesis | 2 | 1 | EC.5.1.3.10 |
| PWY-5835 | geranyl acetate biosynthesis | 2 | 1 | EC.2.3.1.84 |
| PWY-5855 | ubiquinol-7 biosynthesis (prokaryotic) | 6 | 1 | EC.2.1.1.64 |
| PWY-5874 | heme degradation | 2 | 1 | EC.1.14.99.3 |
| PWY-5901 | 2,3-dihydroxybenzoate biosynthesis | 3 | 2 | EC.1.3.1.28 EC.3.3.2.1 |
| PWY-5907 | homospermidine biosynthesis | 1 | 1 | EC.2.5.1.45 |
| PWY-5913 | TCA cycle VI (obligate autotrophs) | 10 | 1 | EC.2.6.1.1 |
| PWY-5921 | glutaminyl-tRNA^gln^ biosynthesis via transamidation | 1 | 1 | EC.6.3.5.7 |
| PWY-5923 | limonene degradation I (D-limonene) | 3 | 2 | EC.1.14.13.107 EC.3.3.2.8 |
| PWY-5929 | puromycin biosynthesis | 1 | 1 | EC.2.1.1.38 |
| PWY-5935 | tuberculosinol biosynthesis | 2 | 1 | EC.5.5.1.16 |
| PWY-5938 | (R)-acetoin biosynthesis I | 2 | 1 | EC.1.1.1.303 |
| PWY-5939 | (R)-acetoin biosynthesis II | 2 | 1 | EC.4.1.1.5 |
| PWY-5940 | streptomycin biosynthesis | 13 | 2 | EC.2.1.4.2 EC.2.6.1.50 |
| PWY-5941 | glycogen degradation II (eukaryotic) | 6 | 4 | EC.2.4.1.1 EC.2.4.1.25 EC.3.2.1.3 EC.3.2.1.33 |
| PWY-5951 | (R,R)-butanediol biosynthesis | 1 | 1 | EC.1.1.1.4 |
| PWY-5958 | acridone alkaloid biosynthesis | 4 | 1 | EC.4.1.3.27 |
| PWY-5964 | guanylyl molybdenum cofactor biosynthesis | 1 | 1 | EC.2.7.7.77 |
| PWY-5965 | fatty acid biosynthesis initiation III | 4 | 1 | EC.2.3.1.41 |
| PWY-5966 | fatty acid biosynthesis initiation II | 4 | 1 | EC.2.3.1.38 |
| PWY-5971 | palmitate biosynthesis II (bacteria and plants) | 9 | 2 | EC.1.3.1.9 EC.4.2.1.59 |
| PWY-5987 | sorgoleone biosynthesis | 6 | 2 | EC.1.2.1.11 EC.1.14.19.1 |
| PWY-5994 | palmitate biosynthesis I (animals and fungi) | 11 | 1 | EC.1.3.1.10 |
| PWY-6000 | γ-linolenate biosynthesis II (animals) | 2 | 1 | EC.1.14.19.3 |
| PWY-6011 | amygdalin and prunasin degradation | 3 | 1 | EC.4.1.2.10 |
| PWY-6012 | acyl carrier protein metabolism I | 2 | 2 | EC.2.7.8.7 EC.3.1.4.14 |
| PWY-6019 | pseudouridine degradation | 2 | 2 | EC.2.7.1.83 EC.4.2.1.70 |
| PWY-6030 | serotonin and melatonin biosynthesis | 4 | 2 | EC.1.14.16.4 EC.2.1.1.4 |
| PWY-6032 | cardenolide biosynthesis | 3 | 1 | EC.1.3.1.22 |
| PWY-6038 | citrate degradation | 2 | 1 | EC.2.8.3.10 |
| PWY-6044 | methanesulfonate degradation | 1 | 1 | EC.1.14.13.111 |
| PWY-6046 | dimethylsulfoniopropanoate degradation I (cleavage) | 1 | 1 | EC.4.4.1.3 |
| PWY-6061 | bile acid biosynthesis, neutral pathway | 12 | 3 | EC.2.3.1.176 EC.4.2.1.107 EC.5.1.99.4 |
| PWY-6073 | alginate biosynthesis I (algal) | 3 | 1 | EC.1.1.1.132 |
| PWY-6074 | zymosterol biosynthesis | 4 | 1 | EC.1.1.1.170 |
| PWY-6077 | anthranilate degradation II (aerobic) | 2 | 2 | EC.1.14.13.40 EC.6.2.1.32 |
| PWY-6087 | 4-chlorocatechol degradation | 3 | 2 | EC.3.1.1.45 EC.5.5.1.7 |
| PWY-6098 | diploterol and cycloartenol biosynthesis | 4 | 1 | EC.5.4.99.17 |
| PWY-6100 | L-carnitine biosynthesis | 3 | 1 | EC.1.14.11.8 |
| PWY-6107 | chlorosalicylate degradation | 3 | 1 | EC.1.14.13.1 |
| PWY-6120 | L-tyrosine biosynthesis III | 3 | 1 | EC.1.3.1.43 |
| PWY-6121 | 5-aminoimidazole ribonucleotide biosynthesis I | 5 | 5 | EC.2.1.2.2 EC.2.4.2.14 EC.6.3.3.1 EC.6.3.4.13 EC.6.3.5.3 |
| PWY-6123 | inosine-5'-phosphate biosynthesis I | 6 | 6 | EC.2.1.2.3 EC.3.5.4.10 EC.4.3.2.2 EC.5.4.99.18 EC.6.3.2.6 EC.6.3.4.18 |
| PWY-6124 | inosine-5'-phosphate biosynthesis II | 5 | 1 | EC.4.1.1.21 |
| PWY-6130 | glycerol degradation III | 2 | 2 | EC.1.1.1.202 EC.4.2.1.30 |
| PWY-6131 | glycerol degradation II | 2 | 1 | EC.1.1.1.6 |
| PWY-6138 | CMP-N-acetylneuraminate biosynthesis I (eukaryotes) | 5 | 4 | EC.2.5.1.57 EC.2.7.1.60 EC.2.7.7.43 EC.3.1.3.29 |
| PWY-6139 | CMP-N-acetylneuraminate biosynthesis II (bacteria) | 3 | 1 | EC.2.5.1.56 |
| PWY-6147 | 6-hydroxymethyl-dihydropterin diphosphate biosynthesis I | 5 | 2 | EC.2.7.6.3 EC.4.1.2.25 |
| PWY-6151 | S-adenosyl-L-methionine cycle I | 4 | 2 | EC.3.2.2.9 EC.4.4.1.21 |
| PWY-6157 | autoinducer AI-1 biosynthesis | 1 | 1 | EC.2.3.1.184 |
| PWY-6163 | chorismate biosynthesis from 3-dehydroquinate | 6 | 5 | EC.1.1.1.282 EC.2.5.1.19 EC.2.7.1.71 EC.4.2.1.10 EC.4.2.3.5 |
| PWY-6164 | 3-dehydroquinate biosynthesis I | 2 | 2 | EC.2.5.1.54 EC.4.2.3.4 |
| PWY-6167 | flavin biosynthesis II (archaea) | 9 | 4 | EC.2.5.1.9 EC.2.7.7.2 EC.3.5.4.29 EC.4.1.99.12 |
| PWY-6168 | flavin biosynthesis III (fungi) | 8 | 1 | EC.3.5.4.25 |
| PWY-6173 | histamine biosynthesis | 1 | 1 | EC.4.1.1.22 |
| PWY-6174 | mevalonate pathway II (archaea) | 7 | 3 | EC.1.1.1.34 EC.2.3.3.10 EC.2.7.1.36 |
| PWY-6181 | histamine degradation | 3 | 1 | EC.2.1.1.8 |
| PWY-6185 | 4-methylcatechol degradation (ortho cleavage) | 6 | 1 | EC.5.3.3.4 |
| PWY-621 | sucrose degradation III (sucrose invertase) | 6 | 1 | EC.3.2.1.26 |
| PWY-6215 | 4-chlorobenzoate degradation | 4 | 1 | EC.1.14.13.2 |
| PWY-622 | starch biosynthesis | 9 | 2 | EC.2.4.1.21 EC.2.7.7.27 |
| PWY-6221 | 2-chlorobenzoate degradation | 1 | 1 | EC.1.14.12.13 |
| PWY-6223 | gentisate degradation I | 3 | 2 | EC.1.13.11.4 EC.5.2.1.4 |
| PWY-6241 | thyroid hormone biosynthesis | 1 | 1 | EC.1.11.1.8 |
| PWY-6260 | thyroid hormone metabolism I (via deiodination) | 2 | 1 | EC.1.97.1.10 |
| PWY-6281 | L-selenocysteine biosynthesis II (archaea and eukaryotes) | 4 | 2 | EC.2.7.9.3 EC.6.1.1.11 |
| PWY-6303 | methyl indole-3-acetate interconversion | 2 | 1 | EC.3.1.1.1 |
| PWY-6307 | L-tryptophan degradation X (mammalian, via tryptamine) | 4 | 1 | EC.1.1.1.2 |
| PWY-6308 | L-cysteine biosynthesis II (tRNA-dependent) | 3 | 1 | EC.3.1.1.29 |
| PWY-6309 | L-tryptophan degradation XI (mammalian, via kynurenine) | 7 | 1 | EC.2.6.1.7 |
| PWY-6317 | galactose degradation I (Leloir pathway) | 5 | 1 | EC.2.7.7.12 |
| PWY-6322 | phosphinothricin tripeptide biosynthesis | 9 | 1 | EC.2.7.8.23 |
| PWY-6344 | L-ornithine degradation II (Stickland reaction) | 7 | 4 | EC.1.4.1.12 EC.5.1.1.12 EC.5.1.1.4 EC.5.4.3.5 |
| PWY-6348 | phosphate acquisition | 1 | 1 | EC.3.1.3.2 |
| PWY-6349 | CDP-archaeol biosynthesis | 5 | 3 | EC.1.1.1.261 EC.2.5.1.41 EC.2.5.1.42 |
| PWY-6351 | D-myo-inositol (1,4,5)-trisphosphate biosynthesis | 5 | 2 | EC.2.7.1.68 EC.3.1.4.11 |
| PWY-6363 | D-myo-inositol (1,4,5)-trisphosphate degradation | 3 | 1 | EC.3.1.3.57 |
| PWY-6373 | acrylate degradation | 3 | 1 | EC.1.2.1.18 |
| PWY-6374 | vibriobactin biosynthesis | 1 | 1 | EC.2.7.7.58 |
| PWY-6386 | UDP-N-acetylmuramoyl-pentapeptide biosynthesis II (lysine-containing) | 8 | 6 | EC.2.5.1.7 EC.5.1.1.3 EC.6.3.2.10 EC.6.3.2.4 EC.6.3.2.8 EC.6.3.2.9 |
| PWY-6387 | UDP-N-acetylmuramoyl-pentapeptide biosynthesis I (meso-diaminopimelate containing) | 8 | 1 | EC.6.3.2.13 |
| PWY-6388 | (S,S)-butanediol degradation | 1 | 1 | EC.1.1.1.76 |
| PWY-6389 | (S)-acetoin biosynthesis | 2 | 1 | EC.1.1.1.304 |
| PWY-6397 | mycolyl-arabinogalactan-peptidoglycan complex biosynthesis | 12 | 1 | EC.5.4.99.9 |
| PWY-6405 | Rapoport-Luebering glycolytic shunt | 3 | 1 | EC.3.1.3.13 |
| PWY-6420 | pyrroloquinoline quinone biosynthesis | 2 | 1 | EC.1.3.3.11 |
| PWY-6421 | arsenate detoxification III (mycothiol) | 3 | 1 | EC.1.8.1.15 |
| PWY-6431 | 4-hydroxybenzoate biosynthesis IV | 2 | 1 | EC.1.2.1.64 |
| PWY-6453 | stigma estolide biosynthesis | 5 | 1 | EC.3.1.3.4 |
| PWY-6464 | polyvinyl alcohol degradation | 1 | 1 | EC.1.1.2.6 |
| PWY-6466 | pyridoxal 5'-phosphate biosynthesis II | 1 | 1 | EC.4.3.3.6 |
| PWY-6476 | cytidylyl molybdenum cofactor biosynthesis | 1 | 1 | EC.2.7.7.76 |
| PWY-6482 | diphthamide biosynthesis (archaea) | 3 | 1 | EC.2.1.1.98 |
| PWY-6483 | ceramide degradation | 1 | 1 | EC.3.5.1.23 |
| PWY-6497 | D-galactarate degradation II | 3 | 1 | EC.4.2.1.42 |
| PWY-6498 | eumelanin biosynthesis | 1 | 1 | EC.5.3.3.12 |
| PWY-6499 | D-glucarate degradation II | 3 | 1 | EC.4.2.1.40 |
| PWY-6507 | 4-deoxy-L-threo-hex-4-enopyranuronate degradation | 6 | 1 | EC.5.3.1.17 |
| PWY-6512 | hydrogen oxidation III (anaerobic, NADP) | 1 | 1 | EC.1.12.1.3 |
| PWY-6517 | N-acetylglucosamine degradation II | 1 | 1 | EC.2.7.1.59 |
| PWY-6518 | glycocholate metabolism (bacteria) | 6 | 2 | EC.1.1.1.159 EC.3.5.1.24 |
| PWY-6519 | 8-amino-7-oxononanoate biosynthesis I | 7 | 1 | EC.2.3.1.47 |
| PWY-6527 | stachyose degradation | 8 | 1 | EC.3.2.1.22 |
| PWY-6529 | chlorate reduction | 2 | 1 | EC.1.13.11.49 |
| PWY-6531 | mannitol cycle | 4 | 2 | EC.1.1.1.17 EC.1.1.1.67 |
| PWY-6536 | 4-aminobutanoate degradation III | 2 | 1 | EC.1.2.1.16 |
| PWY-6538 | caffeine degradation III (bacteria, via demethylation) | 5 | 1 | EC.1.17.3.2 |
| PWY-6543 | 4-aminobenzoate biosynthesis | 2 | 2 | EC.2.6.1.85 EC.4.1.3.38 |
| PWY-6545 | pyrimidine deoxyribonucleotides de novo biosynthesis III | 9 | 7 | EC.1.17.4.1 EC.2.1.1.148 EC.2.7.4.13 EC.2.7.4.9 EC.3.6.1.15 EC.3.6.1.19 EC.3.6.1.23 |
| PWY-6550 | carbazole degradation | 3 | 1 | EC.1.13.11.39 |
| PWY-6556 | pyrimidine ribonucleosides salvage II | 2 | 2 | EC.3.2.2.3 EC.3.5.4.5 |
| PWY-6558 | heparan sulfate biosynthesis (late stages) | 10 | 1 | EC.2.8.2.23 |
| PWY-6562 | norspermidine biosynthesis | 6 | 1 | EC.4.1.1.86 |
| PWY-6572 | chondroitin sulfate degradation I (bacterial) | 7 | 2 | EC.4.2.2.20 EC.4.2.2.5 |
| PWY-6578 | 8-amino-7-oxononanoate biosynthesis III | 2 | 1 | EC.6.2.1.14 |
| PWY-6596 | adenosine nucleotides degradation I | 7 | 1 | EC.3.2.2.1 |
| PWY-6599 | guanine and guanosine salvage II | 2 | 1 | EC.2.4.2.8 |
| PWY-66 | GDP-L-fucose biosynthesis I (from GDP-D-mannose) | 2 | 1 | EC.1.1.1.271 |
| PWY-6609 | adenine and adenosine salvage III | 3 | 1 | EC.3.5.4.4 |
| PWY-6610 | adenine and adenosine salvage IV | 3 | 1 | EC.3.5.4.2 |
| PWY-6611 | adenine and adenosine salvage V | 3 | 1 | EC.2.7.1.73 |
| PWY-6614 | tetrahydrofolate biosynthesis | 3 | 2 | EC.2.5.1.15 EC.6.3.2.12 |
| PWY-6616 | sulfolactate degradation I | 3 | 1 | EC.4.4.1.24 |
| PWY-6617 | adenosine nucleotides degradation III | 1 | 1 | EC.3.2.2.4 |
| PWY-6619 | adenine and adenosine salvage VI | 1 | 1 | EC.2.7.1.20 |
| PWY-6622 | heptadecane biosynthesis | 3 | 1 | EC.6.2.1.20 |
| PWY-6638 | sulfolactate degradation III | 3 | 1 | EC.4.4.1.25 |
| PWY-6646 | fluoroacetate degradation | 1 | 1 | EC.3.8.1.3 |
| PWY-6649 | glycolate and glyoxylate degradation III | 3 | 1 | EC.1.1.99.14 |
| PWY-6672 | cis-genanyl-CoA degradation | 8 | 2 | EC.6.2.1.1 EC.6.4.1.5 |
| PWY-6683 | sulfate reduction III (assimilatory) | 3 | 2 | EC.1.8.1.2 EC.1.8.4.10 |
| PWY-6690 | cinnamate and 3-hydroxycinnamate degradation to 2-oxopent-4-enoate | 5 | 2 | EC.1.13.11.16 EC.1.14.12.19 |
| PWY-6695 | oxalate degradation II | 3 | 2 | EC.2.8.3.16 EC.4.1.1.8 |
| PWY-6698 | oxalate degradation V | 1 | 1 | EC.4.1.1.2 |
| PWY-6700 | queuosine biosynthesis | 4 | 2 | EC.1.7.1.13 EC.2.4.2.29 |
| PWY-6713 | L-rhamnose degradation II | 7 | 1 | EC.4.2.1.90 |
| PWY-6724 | starch degradation II | 6 | 1 | EC.3.2.1.2 |
| PWY-6731 | starch degradation III | 6 | 1 | EC.3.2.1.54 |
| PWY-6745 | phytochelatins biosynthesis | 1 | 1 | EC.2.3.2.15 |
| PWY-6748 | nitrate reduction VII (denitrification) | 4 | 1 | EC.1.7.2.4 |
| PWY-6749 | CMP-legionaminate biosynthesis I | 9 | 2 | EC.2.6.1.16 EC.5.4.2.10 |
| PWY-6754 | S-methyl-5'-thioadenosine degradation I | 2 | 2 | EC.2.7.1.100 EC.3.2.2.16 |
| PWY-6756 | S-methyl-5'-thioadenosine degradation II | 1 | 1 | EC.2.4.2.28 |
| PWY-6759 | hydrogen production III | 1 | 1 | EC.1.12.7.2 |
| PWY-6784 | cellulose and hemicellulose degradation (cellulolosome) | 3 | 2 | EC.3.1.1.73 EC.3.2.1.8 |
| PWY-6785 | hydrogen production VIII | 5 | 1 | EC.1.6.5.2 |
| PWY-6788 | cellulose degradation II (fungi) | 3 | 2 | EC.3.2.1.4 EC.3.2.1.91 |
| PWY-6789 | (1,3)-β-D-xylan degradation | 2 | 1 | EC.3.2.1.32 |
| PWY-6803 | phosphatidylcholine acyl editing | 4 | 2 | EC.3.1.1.32 EC.3.1.1.4 |
| PWY-6807 | xyloglucan degradation II (exoglucanase) | 4 | 1 | EC.3.2.1.23 |
| PWY-681 | dibenzothiophene desulfurization | 3 | 1 | EC.3.13.1.3 |
| PWY-6815 | porphyran degradation | 3 | 2 | EC.3.2.1.159 EC.3.2.1.81 |
| PWY-6821 | κ-carrageenan degradation | 3 | 1 | EC.3.2.1.83 |
| PWY-6822 | ι-carrageenan degradation | 1 | 1 | EC.3.2.1.157 |
| PWY-6823 | molybdenum cofactor biosynthesis | 7 | 5 | EC.2.10.1.1 EC.2.7.7.75 EC.2.8.1.12 EC.2.8.1.7 EC.4.1.99.18 |
| PWY-6825 | phosphatidylcholine biosynthesis V | 2 | 1 | EC.2.1.1.17 |
| PWY-6826 | phosphatidylcholine biosynthesis VI | 1 | 1 | EC.2.7.8.24 |
| PWY-6829 | tRNA methylation (yeast) | 10 | 1 | EC.2.1.1.228 |
| PWY-6834 | spermidine biosynthesis III | 4 | 1 | EC.4.1.1.50 |
| PWY-6840 | homoglutathione biosynthesis | 2 | 1 | EC.6.3.2.2 |
| PWY-6848 | rutin degradation | 3 | 1 | EC.1.13.11.24 |
| PWY-6854 | ethylene biosynthesis III (microbes) | 3 | 1 | EC.1.15.1.1 |
| PWY-6857 | retinol biosynthesis | 5 | 2 | EC.1.14.99.36 EC.3.1.1.3 |
| PWY-6863 | pyruvate fermentation to hexanol | 9 | 1 | EC.1.3.1.44 |
| PWY-6871 | 3-methylbutanol biosynthesis | 3 | 1 | EC.2.3.3.13 |
| PWY-6872 | retinoate biosynthesis I | 2 | 1 | EC.1.1.1.105 |
| PWY-6890 | 4-amino-2-methyl-5-diphosphomethylpyrimidine biosynthesis | 2 | 1 | EC.2.7.4.7 |
| PWY-6891 | thiazole biosynthesis II (Bacillus) | 5 | 1 | EC.2.2.1.7 |
| PWY-6892 | thiazole biosynthesis I (E. coli) | 6 | 1 | EC.4.1.99.19 |
| PWY-6893 | thiamin diphosphate biosynthesis II (Bacillus) | 2 | 2 | EC.2.5.1.3 EC.2.7.4.16 |
| PWY-6897 | thiamin salvage II | 3 | 1 | EC.2.7.1.50 |
| PWY-6898 | thiamin salvage III | 1 | 1 | EC.2.7.6.2 |
| PWY-6899 | base-degraded thiamin salvage | 2 | 1 | EC.3.5.99.2 |
| PWY-6902 | chitin degradation II | 2 | 2 | EC.3.2.1.14 EC.3.2.1.52 |
| PWY-6906 | chitin derivatives degradation | 7 | 1 | EC.3.5.1.25 |
| PWY-6910 | hydroxymethylpyrimidine salvage | 2 | 1 | EC.2.7.1.49 |
| PWY-6915 | pentalenolactone biosynthesis | 7 | 1 | EC.4.2.3.7 |
| PWY-6936 | seleno-amino acid biosynthesis | 5 | 3 | EC.2.3.1.30 EC.2.5.1.47 EC.4.4.1.8 |
| PWY-6943 | testosterone and androsterone degradation to androstendione | 5 | 1 | EC.1.3.99.5 |
| PWY-6944 | androstenedione degradation | 14 | 3 | EC.1.1.1.145 EC.1.14.13.142 EC.1.3.99.4 |
| PWY-6945 | cholesterol degradation to androstenedione I (cholesterol oxidase) | 12 | 1 | EC.1.1.3.6 |
| PWY-695 | abscisic acid biosynthesis | 5 | 1 | EC.1.13.11.51 |
| PWY-6952 | glycerophosphodiester degradation | 2 | 1 | EC.3.1.4.46 |
| PWY-6961 | L-ascorbate degradation II (bacterial, aerobic) | 5 | 4 | EC.1.1.1.130 EC.4.1.1.85 EC.5.1.3.22 EC.5.1.3.4 |
| PWY-6963 | ammonia assimilation cycle I | 2 | 1 | EC.1.4.1.14 |
| PWY-6965 | methylamine degradation II | 3 | 2 | EC.1.5.99.5 EC.2.1.1.21 |
| PWY-6966 | methanol oxidation to formaldehyde I | 1 | 1 | EC.1.1.2.7 |
| PWY-6967 | methylamine degradation I | 1 | 1 | EC.1.4.9.1 |
| PWY-6970 | acetyl-CoA biosynthesis II (NADP-dependent pyruvate dehydrogenase) | 1 | 1 | EC.1.2.1.51 |
| PWY-6986 | alginate degradation | 6 | 1 | EC.4.2.2.3 |
| PWY-6987 | lipoate biosynthesis and incorporation III (Bacillus) | 3 | 2 | EC.2.3.1.181 EC.2.8.1.8 |
| PWY-6992 | 1,5-anhydrofructose degradation | 5 | 1 | EC.1.1.1.292 |
| PWY-7002 | 4-hydroxyacetophenone degradation | 5 | 1 | EC.1.14.13.84 |
| PWY-7007 | methyl ketone biosynthesis | 6 | 1 | EC.6.2.1.2 |
| PWY-7013 | L-1,2-propanediol degradation | 5 | 1 | EC.4.2.1.28 |
| PWY-702 | L-methionine biosynthesis II | 5 | 1 | EC.2.7.1.39 |
| PWY-7039 | phosphatidate metabolism, as a signaling molecule | 5 | 1 | EC.2.7.1.107 |
| PWY-7052 | cyanophycin metabolism | 3 | 2 | EC.6.3.2.29 EC.6.3.2.30 |
| PWY-7077 | N-acetyl-D-galactosamine degradation | 5 | 2 | EC.2.7.1.144 EC.4.1.2.40 |
| PWY-7084 | nitrifier denitrification | 4 | 1 | EC.1.7.2.5 |
| PWY-7090 | UDP-2,3-diacetamido-2,3-dideoxy-α-D-mannuronate biosynthesis | 5 | 1 | EC.1.1.1.136 |
| PWY-7098 | vanillin and vanillate degradation II | 2 | 1 | EC.1.14.13.82 |
| PWY-7106 | erythromycin D biosynthesis | 4 | 1 | EC.2.3.1.94 |
| PWY-7115 | C4 photosynthetic carbon assimilation cycle, NAD-ME type | 8 | 1 | EC.2.6.1.2 |
| PWY-7118 | chitin degradation to ethanol | 7 | 1 | EC.3.5.1.41 |
| PWY-7119 | sphingolipid recycling and degradation (yeast) | 5 | 1 | EC.4.1.2.27 |
| PWY-7130 | L-glucose degradation | 6 | 2 | EC.2.7.1.58 EC.4.1.2.21 |
| PWY-7134 | rutin degradation (plants) | 2 | 1 | EC.3.2.1.40 |
| PWY-7139 | sesaminol glucoside biosynthesis | 2 | 1 | EC.1.2.1.11 |
| PWY-7142 | cyanide detoxification II | 2 | 1 | EC.3.5.1.49 |
| PWY-7158 | L-phenylalanine degradation V | 3 | 1 | EC.1.5.1.34 |
| PWY-7165 | L-ascorbate biosynthesis VI (engineered pathway) | 6 | 3 | EC.1.1.1.274 EC.1.1.5.2 EC.1.1.99.3 |
| PWY-7176 | UTP and CTP de novo biosynthesis | 4 | 3 | EC.2.7.4.14 EC.2.7.4.22 EC.6.3.4.2 |
| PWY-7180 | 2'-deoxy-α-D-ribose 1-phosphate degradation | 3 | 2 | EC.4.1.2.4 EC.5.4.2.7 |
| PWY-7181 | pyrimidine deoxyribonucleosides degradation | 4 | 3 | EC.2.4.2.2 EC.2.4.2.3 EC.2.4.2.4 |
| PWY-7183 | pyrimidine nucleobases salvage I | 1 | 1 | EC.2.4.2.9 |
| PWY-7187 | pyrimidine deoxyribonucleotides de novo biosynthesis II | 9 | 2 | EC.1.17.4.2 EC.3.5.4.13 |
| PWY-7193 | pyrimidine ribonucleosides salvage I | 2 | 1 | EC.2.7.1.48 |
| PWY-7194 | pyrimidine nucleobases salvage II | 1 | 1 | EC.3.5.4.1 |
| PWY-7199 | pyrimidine deoxyribonucleosides salvage | 5 | 2 | EC.2.7.1.21 EC.2.7.1.74 |
| PWY-7204 | pyridoxal 5'-phosphate salvage II (plants) | 4 | 2 | EC.1.1.1.65 EC.3.1.3.74 |
| PWY-7206 | pyrimidine deoxyribonucleotides dephosphorylation | 5 | 1 | EC.3.6.1.12 |
| PWY-7210 | pyrimidine deoxyribonucleotides biosynthesis from CTP | 9 | 1 | EC.3.5.4.12 |
| PWY-7219 | adenosine ribonucleotides de novo biosynthesis | 4 | 2 | EC.2.7.4.3 EC.6.3.4.4 |
| PWY-7221 | guanosine ribonucleotides de novo biosynthesis | 4 | 2 | EC.2.7.4.8 EC.6.3.5.2 |
| PWY-7224 | purine deoxyribonucleosides salvage | 8 | 2 | EC.2.7.1.113 EC.2.7.1.76 |
| PWY-723 | alkylnitronates degradation | 2 | 1 | EC.1.13.12.16 |
| PWY-7238 | sucrose biosynthesis II | 11 | 2 | EC.2.4.1.14 EC.3.1.3.24 |
| PWY-7242 | D-fructuronate degradation | 6 | 2 | EC.1.1.1.57 EC.4.2.1.8 |
| PWY-7247 | β-D-glucuronide and D-glucuronate degradation | 2 | 2 | EC.3.2.1.31 EC.5.3.1.12 |
| PWY-7254 | TCA cycle VII (acetate-producers) | 7 | 1 | EC.1.1.5.4 |
| PWY-7255 | ergothioneine biosynthesis I (bacteria) | 4 | 1 | EC.2.1.1.44 |
| PWY-7270 | L-methionine salvage cycle II (plants) | 1 | 1 | EC.4.4.1.14 |
| PWY-7277 | sphingolipid biosynthesis (mammals) | 3 | 1 | EC.3.1.4.12 |
| PWY-7290 | Escherichia coli serotype O86 O-antigen biosynthesis | 6 | 1 | EC.2.7.8.33 |
| PWY-7294 | xylose degradation IV | 7 | 1 | EC.1.1.1.26 |
| PWY-7301 | dTDP-β-L-noviose biosynthesis | 5 | 1 | EC.5.1.3.13 |
| PWY-7303 | 3-dimethylallyl-4-hydroxybenzoate biosynthesis | 4 | 1 | EC.1.3.1.12 |
| PWY-7309 | acrylonitrile degradation II | 1 | 1 | EC.3.5.5.7 |
| PWY-7310 | D-glucosaminate degradation | 4 | 1 | EC.2.7.1.69 |
| PWY-7312 | dTDP-D-β-fucofuranose biosynthesis | 4 | 1 | EC.1.1.1.266 |
| PWY-7335 | UDP-N-acetyl-α-D-mannosaminouronate biosynthesis | 2 | 2 | EC.1.1.1.336 EC.5.1.3.14 |
| PWY-7346 | UDP-α-D-glucuronate biosynthesis (from UDP-glucose) | 1 | 1 | EC.1.1.1.22 |
| PWY-735 | jasmonic acid biosynthesis | 10 | 3 | EC.1.3.1.42 EC.3.1.2.20 EC.4.2.1.92 |
| PWY-7375 | mRNA capping I | 3 | 2 | EC.2.1.1.56 EC.2.7.7.50 |
| PWY-7376 | cob(II)yrinate a,c-diamide biosynthesis II (late cobalt incorporation) | 9 | 7 | EC.1.3.1.54 EC.2.1.1.130 EC.2.1.1.131 EC.2.1.1.133 EC.2.1.1.152 EC.6.3.5.9 EC.6.6.1.2 |
| PWY-7377 | cob(II)yrinate a,c-diamide biosynthesis I (early cobalt insertion) | 11 | 2 | EC.2.1.1.151 EC.4.99.1.3 |
| PWY-7378 | aminopropanol phosphate biosynthesis II | 2 | 1 | EC.1.1.1.103 |
| PWY-7380 | biotin biosynthesis from 8-amino-7-oxononanoate II | 3 | 2 | EC.2.8.1.6 EC.6.3.3.3 |
| PWY-7383 | anaerobic energy metabolism (invertebrates, cytosol) | 6 | 2 | EC.4.1.1.32 EC.5.1.1.1 |
| PWY-7391 | isoprene biosynthesis II (engineered) | 8 | 2 | EC.2.7.4.2 EC.4.1.1.33 |
| PWY-7396 | butanol and isobutanol biosynthesis (engineered) | 7 | 2 | EC.1.1.1.85 EC.1.4.3.19 |
| PWY-7401 | crotonate fermentation (to acetate and cyclohexane carboxylate) | 13 | 1 | EC.4.2.1.100 |
| PWY-7409 | phospholipid remodeling (phosphatidylethanolamine, yeast) | 5 | 1 | EC.3.1.1.5 |
| PWY-7415 | tylosin biosynthesis | 9 | 1 | EC.2.1.1.101 |
| PWY-7416 | phospholipid remodeling (phosphatidylcholine, yeast) | 3 | 1 | EC.2.3.1.158 |
| PWY-7420 | monoacylglycerol metabolism (yeast) | 3 | 1 | EC.3.1.1.23 |
| PWY-7425 | 2-chloroacrylate degradation I | 3 | 1 | EC.3.8.1.2 |
| PWY-7426 | mannosyl-glycoprotein N-acetylglucosaminyltransferases | 7 | 2 | EC.2.4.1.144 EC.3.2.1.114 |
| PWY-7429 | arsenite oxidation II (respiratory) | 1 | 1 | EC.1.20.9.1 |
| PWY-7431 | aromatic biogenic amine degradation (bacteria) | 7 | 1 | EC.1.14.14.9 |
| PWY-7433 | mucin core 1 and core 2 O-glycosylation | 5 | 1 | EC.2.4.1.41 |
| PWY-7456 | mannan degradation | 7 | 2 | EC.3.2.1.78 EC.5.1.3.11 |
| PWY-7459 | kojibiose degradation | 2 | 1 | EC.2.4.1.230 |
| PWY-7515 | trans-3-hydroxy-L-proline degradation | 3 | 1 | EC.4.2.1.77 |
| PWY-7560 | methylerythritol phosphate pathway II | 8 | 6 | EC.1.1.1.267 EC.1.17.1.2 EC.1.17.7.1 EC.2.7.1.148 EC.2.7.7.60 EC.4.6.1.12 |
| PWY-7571 | ferrichrome A biosynthesis | 5 | 1 | EC.4.2.1.18 |
| PWY-7581 | N-acetylneuraminate and N-acetylmannosamine degradation II | 2 | 2 | EC.4.1.3.3 EC.5.1.3.8 |
| PWY-7644 | heparin degradation | 5 | 3 | EC.3.1.6.13 EC.3.10.1.1 EC.4.2.2.7 |
| PWY-7645 | hyaluronan degradation | 3 | 1 | EC.4.2.2.1 |
| PWY-7651 | heparan sulfate degradation | 3 | 1 | EC.4.2.2.8 |
| PWY-7661 | protein N-glycosylation (Haloferax volcanii) | 4 | 1 | EC.2.4.1.83 |
| PWY-7664 | oleate biosynthesis IV (anaerobic) | 7 | 1 | EC.5.3.3.14 |
| PWY-801 | L-homocysteine and L-cysteine interconversion | 1 | 1 | EC.4.2.1.22 |
| PWY-81 | toluene degradation to benzoyl-CoA (anaerobic) | 3 | 2 | EC.2.8.3.15 EC.4.1.99.11 |
| PWY-822 | fructan biosynthesis | 4 | 1 | EC.2.4.1.10 |
| PWY-842 | starch degradation I | 3 | 1 | EC.3.2.1.20 |
| PWY-862 | fructan degradation | 2 | 1 | EC.3.2.1.80 |
| PWY-881 | trehalose biosynthesis II | 2 | 1 | EC.3.1.3.12 |
| PWY0-1182 | trehalose degradation II (trehalase) | 3 | 1 | EC.3.2.1.28 |
| PWY0-1221 | putrescine degradation II | 3 | 2 | EC.3.5.1.94 EC.6.3.1.11 |
| PWY0-1241 | ADP-L-glycero-β-D-manno-heptose biosynthesis | 5 | 1 | EC.5.1.3.20 |
| PWY0-1261 | anhydromuropeptides recycling | 10 | 2 | EC.2.3.1.157 EC.3.5.1.28 |
| PWY0-1264 | biotin-carboxyl carrier protein assembly | 3 | 2 | EC.6.3.4.14 EC.6.3.4.15 |
| PWY0-1309 | chitobiose degradation | 2 | 1 | EC.3.2.1.86 |
| PWY0-1314 | fructose degradation | 1 | 1 | EC.2.7.1.56 |
| PWY0-1324 | N-acetylneuraminate and N-acetylmannosamine degradation I | 3 | 1 | EC.5.1.3.9 |
| PWY0-1347 | NADH to trimethylamine N-oxide electron transfer | 2 | 1 | EC.1.7.2.3 |
| PWY0-1415 | superpathway of heme biosynthesis from uroporphyrinogen-III | 4 | 1 | EC.4.99.1.1 |
| PWY0-1465 | D-malate degradation | 1 | 1 | EC.1.1.1.83 |
| PWY0-1477 | ethanolamine utilization | 1 | 1 | EC.4.3.1.7 |
| PWY0-1479 | tRNA processing | 5 | 4 | EC.2.7.7.56 EC.3.1.13.1 EC.3.1.13.5 EC.3.1.26.5 |
| PWY0-1507 | biotin biosynthesis from 8-amino-7-oxononanoate I | 3 | 1 | EC.2.6.1.62 |
| PWY0-1584 | nitrate reduction X (periplasmic, dissimilatory) | 2 | 1 | EC.1.7.99.4 |
| PWY0-321 | phenylacetate degradation I (aerobic) | 9 | 1 | EC.1.14.13.149 |
| PWY0-41 | allantoin degradation IV (anaerobic) | 4 | 1 | EC.2.7.2.2 |
| PWY0-42 | 2-methylcitrate cycle I | 5 | 1 | EC.4.2.1.79 |
| PWY0-43 | conversion of succinate to propanoate | 3 | 1 | EC.4.1.1.41 |
| PWY0-662 | PRPP biosynthesis I | 1 | 1 | EC.2.7.6.1 |
| PWY0-901 | L-selenocysteine biosynthesis I (bacteria) | 3 | 1 | EC.2.9.1.1 |
| PWY0-981 | taurine degradation IV | 1 | 1 | EC.1.14.11.17 |
| PWY1-2 | L-alanine degradation IV | 1 | 1 | EC.1.4.1.1 |
| PWY1G-0 | mycothiol biosynthesis | 5 | 1 | EC.3.5.1.103 |
| PWY3O-4106 | NAD salvage pathway III | 2 | 2 | EC.2.7.1.22 EC.2.7.7.1 |
| PWY3O-6 | dehydro-D-arabinono-1,4-lactone biosynthesis | 3 | 2 | EC.1.1.1.116 EC.1.1.3.37 |
| PWY490-3 | nitrate reduction VI (assimilatory) | 4 | 1 | EC.1.7.7.2 |
| PWY490-4 | L-asparagine biosynthesis III (tRNA-dependent) | 3 | 1 | EC.6.3.5.6 |
| PWY5F9-12 | biphenyl degradation | 4 | 2 | EC.1.14.12.18 EC.1.3.1.56 |
| PWY5F9-3233 | phthalate degradation | 3 | 1 | EC.4.1.1.69 |
| PWY66-201 | nicotine degradation IV | 9 | 1 | EC.1.14.13.8 |
| PWY66-367 | ketogenesis | 5 | 1 | EC.1.1.1.30 |
| PWY66-368 | ketolysis | 3 | 1 | EC.2.8.3.5 |
| PWY66-373 | sucrose degradation V (sucrose α-glucosidase) | 6 | 2 | EC.2.7.1.3 EC.3.2.1.48 |
| PWY66-375 | leukotriene biosynthesis | 4 | 1 | EC.3.3.2.6 |
| PWY66-380 | estradiol biosynthesis I (via estrone) | 1 | 1 | EC.1.1.1.62 |
| PWY66-389 | phytol degradation | 4 | 1 | EC.1.3.1.38 |
| PWY66-399 | gluconeogenesis III | 13 | 1 | EC.3.1.3.58 |
| PWY66-423 | fructose 2,6-bisphosphate biosynthesis | 2 | 1 | EC.3.1.3.46 |
| PWY66-425 | L-lysine degradation II (L-pipecolate pathway) | 5 | 1 | EC.1.2.1.31 |
| PWY66-428 | L-threonine degradation V | 1 | 1 | EC.4.3.1.19 |
| PYRIDNUCSAL-PWY | NAD salvage pathway I | 7 | 1 | EC.3.5.1.42 |
| PYRIDNUCSYN-PWY | NAD biosynthesis I (from aspartate) | 6 | 1 | EC.6.3.1.5 |
| PYRIDOXSYN-PWY | pyridoxal 5'-phosphate biosynthesis I | 6 | 4 | EC.1.1.1.290 EC.1.2.1.72 EC.2.6.1.52 EC.2.6.99.2 |
| PYRUVDEHYD-PWY | pyruvate decarboxylation to acetyl CoA | 3 | 2 | EC.1.2.4.1 EC.2.3.1.12 |
